# Biochemically validated structural model of the 15-subunit IFT-B complex

**DOI:** 10.1101/2022.08.20.504624

**Authors:** Narcis A. Petriman, Marta Loureiro-López, Michael Taschner, Nevin K. Zacharia, Magdalena M. Georgieva, Niels Boegholm, André Mourão, Robert B. Russell, Jens S. Andersen, Esben Lorentzen

## Abstract

Cilia are ubiquitous eukaryotic organelles important to cellular motility, signalling and sensory reception. Cilium formation requires intraflagellar transport for trafficking of structural and signalling components. The large MDa IFT-B complex constitutes the backbone of polymeric IFT trains that carry ciliary cargo between the cilium and the cell body. Currently, high-resolution structures are only available for smaller IFT-B sub-complexes leaving >50% of the IFT-B complex structurally uncharacterized. We have used recent advances in protein structure prediction as implemented in Alphafold to assemble a structural model for the 15-subunit IFT-B complex. The model was validated using crosslinking/MS data on reconstituted IFT-B complexes, X-ray scattering in solution and diffraction from crystals as well as site-directed mutagenesis and protein binding assays. The IFT-B structural model reveals an elongated and highly flexible complex consistent with cryo-electron tomographic reconstructions of IFT trains. The >400Å long IFT-B complex can roughly be divided into IFT-B1 and IFT-B2 parts with binding sites for ciliary cargo and the inactive IFT dynein motor, respectively. Interestingly, our structural modelling and crosslinking/MS results are consistent with two different binding sites for IFT81/74 on IFT88/70/52/46 suggesting the possibility of two different structural architectures for the IFT-B1 complex. Our data present a structural framework to understand IFT-B complex assembly, function, and ciliopathy variants.

## Introduction

Cilia are slim eukaryotic organelles that are conserved from the green alga *Chlamydomonas reinhardtii* (Cr) to human and protrude from cell surfaces to function in both motility and signalling pathways (Rosenbaum and Witman, 2002). Cilia are organized into an axoneme consisting of microtubule (MT)-doublets with nine-fold symmetry and are surrounded by the ciliary membrane, which is continuous with the plasma membrane but contains a unique composition of lipids and membrane receptors important for signalling (Mourão et al., 2016). Cilium formation and function require the selective ciliary trafficking of both axonemal factors such as tubulin (Bhogaraju et al., 2014) as well as membrane proteins (Long and Huang, 2020). Trafficking along the ciliary axoneme is carried out by intraflagellar transport (IFT) (Kozminski et al., 1993), which relies on molecular motors and the 22 subunit IFT complex that organizes into 6 subunit IFT-A and 16 subunit IFT-B complexes that loosely associate (Cole et al., 1998; Piperno et al., 1998). IFT-A and -B polymerize into linear assemblies known as IFT trains that move ciliary cargo into and out of cilia (Kozminski et al., 1995; Pigino et al., 2009). Anterograde IFT trains move from the base to the tip of cilia powered by the kinesin 2 motor (Kozminski et al., 1995; Wingfield et al., 2017) whereas retrograde IFT trains move from the tip and back to the base of cilia and are powered by the IFT dynein motor (Pazour et al., 1999; Porter et al., 1999). Elegant time-resolved correlative fluorescence and three-dimensional electron microscopy revealed that anterograde and retrograde IFT trains drive on different MTs of the MT doublets to avoid head-on collisions (Stepanek and Pigino, 2016). During kinesin-driven IFT to the ciliary tip, inactivated IFT dynein motor associate with anterograde IFT trains as a cargo (Jordan et al., 2018).

Interestingly, IFT-B and IFT-A assemble at the ciliary base into linear polymers of different repeat length (Jordan et al., 2018; van den Hoek et al., 2022). Whereas IFT-B polymers have a repeat distance of 6nm and form first, IFT-A has a repeat distance of 11.5nm and appear to assemble onto preformed IFT-B polymers (Jordan et al., 2018; van den Hoek et al., 2022). IFT-A and IFT-B thus do not form 1:1 complexes but rather have an approximate 1:2 stoichiometry in IFT trains, which is consistent with mass-spectrometry (MS) results (Lechtreck et al., 2009). The structures of anterograde IFT trains were determined at 24-37Å resolution by cryo-electron tomography (cryo-ET), which clearly resolved IFT-A, IFT-B and inactive IFT dynein complexes (Jordan et al., 2018; van den Hoek et al., 2022). However, the resolution of these studies was insufficient to resolve the position of individual protein subunits. Interestingly, retrograde IFT trains returning from the ciliary tip to the base appear to have very different structures and repeat distances compared to anterograde IFT trains suggesting that significant remodelling of the IFT complexes occurs at or close to the ciliary tip (Jordan et al., 2018).

The IFT-B complex forms the backbone of IFT trains and is absolutely required for IFT and thus for cilium formation (Taschner and Lorentzen, 2016). Biochemical studies have provided an architecture of the IFT-B complex (Boldt et al., 2016; Katoh et al., 2016; Taschner et al., 2016), sometimes with domain resolution, and several high-resolution crystal structures have been determined for IFT subunits and smaller sub-complexes (Figure 1A) (Taschner and Lorentzen, 2016). These include the structure of the IFT27/25 complex (Bhogaraju et al., 2011) involved in BBSome trafficking and hedgehog signalling (Desai et al., 2020; Eguether et al., 2014; Keady et al., 2012), IFT81N/74N/22 revealing the binding mode of the small GTPase IFT22 on IFT81/74 (Wachter et al., 2019), and the IFT70/52 and IFT52/46 sub-complexes demonstrating how IFT70 wraps around IFT52 as a superhelix (Taschner et al., 2014). The crystal structure of IFT80 revealed the structure of two β-propellers (BP), suggested an IFT80 homo-dimer, and allowed for the mechanistic study of ciliopathy disease mutations (Taschner et al., 2018). In addition, crystal structures are available for the N-terminal IFT54 calponin homology (CH) and IFT52 GIFT domains (Taschner et al., 2016). These studies have established how the IFT-B complex organizes into approximately equally sized IFT-B1 and IFT-B2 complexes that associate via IFT88/52 of IFT-B1 and IFT57/38 of IFT-B2 (Katoh et al., 2016; Taschner et al., 2016). However, the high-resolution structures of IFT proteins cannot be unambiguously fitted to the low resolution cryo-ET maps thus preventing a structural understanding of the complete IFT-B complex.

**Figure 1:**
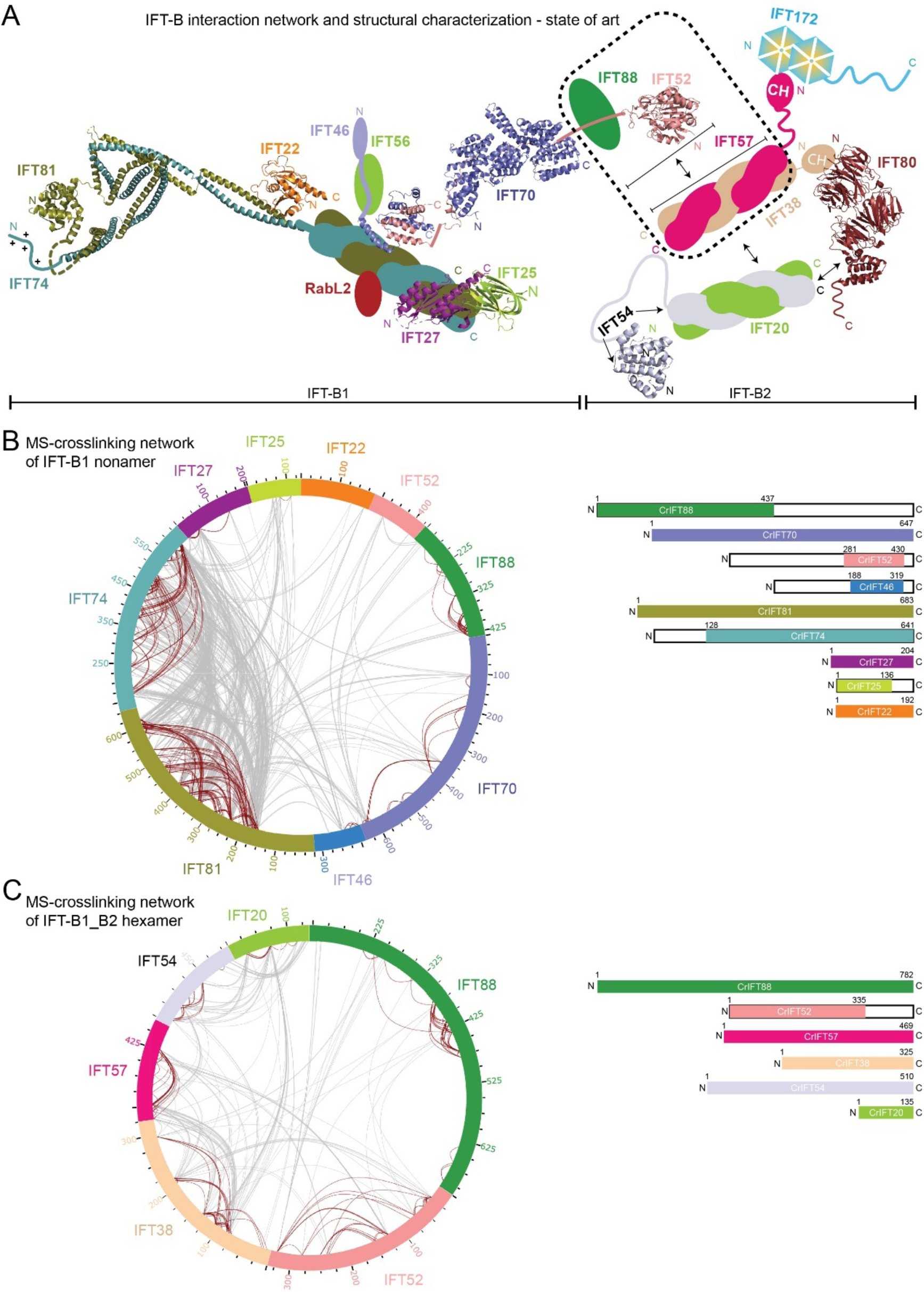
Interaction network of the IFT-B complex obtained by chemical cross-linking and mass spectrometry. **(A)** Schematic representation of IFT-B complex architecture based on published structural and biochemical data. Structural information is available for *Trypanosoma brucei* IFT22 and the N-termini of IFT81/74 (PDB:6ian), the *Chlamydomonas reinhardtii* IFT27/25 heterodimer (PDB: 2yc2), the C-termini of IFT52 and IFT46 from *Tetrahymena thermophila* (PDB: 4uzz), the IFT70/52 complex from *Chlamydomonas reinhardtii* (4uzy), the globular N-terminal GIFT domain of IFT52 from *Chlamydomonas reinhardtii* (PDB: 5FMR) as well as from *Mus musculus* (PDB: 5FMS), the N-terminal CH-domain of IFT54 from *Mus musculus* (5FMU) and IFT80 from *Chamydomonas reinhardtii* (PDB: 5N4A). The IFT-B complex is subdivided into biochemically salt stable IFT-B1 (IFT88/81/74/70/56/52/46/27/25/22/RabL2) and IFT-B2 (IFT172/80/57/54/38/20) complexes. **(B**) The inter- and intra-molecular crosslinking network within the *Chlamydomonas* IFT-B1 nonamer (IFT88_1-437_/70/52_281-430_/46_188-319_/81/74_128-C_/27/25_1-136_/22) are depicted as a cartwheel diagram (left panel). The grey lines show intermolecular crosslinking pairs, and the brown lines show the intramolecular crosslinking pairs. The protein constructs of the IFT-B1 nonamer are indicated on the right. **(C)** The inter- and intra-molecular crosslinking network within a CrIFT-B1-B2 hexamer (IFT88/52_1-335_/57/38/54/20) is displayed as a cartwheel. In this protein complex only the C-terminal part of IFT52 is truncated while all other proteins are full length (see schematics on the right panel).

To bridge this gap in our understanding of the IFT-B structure, we use recent advances in machine learning to model the structure of IFT-B sub-complexes and assemble an almost complete 15-subunit structural model of the IFT-B complex. We use a plethora of biochemical and biophysical methods to validate interactions and interfaces within the IFT-B complex. Our structural model is consistent with cryo-ET maps and provide a structural framework to understand IFT and allows for the mapping of ciliopathy variants in context of the IFT-B complex.

## Results

### Crosslinking/MS reveals the interactions within the IFT-B complex

To obtain a comprehensive map of interactions within the IFT-B complex, we produced two recombinant *Chlamydomonas* IFT-B complexes covering 13 subunits and analysed these by crosslinking/mass spectrometry (MS). We enriched an IFT-B1 complex consisting of the nine *Chlamydomonas* proteins IFT88, IFT81, IFT74, IFT70, IFT52, IFT46, IFT27, IFT25 and IFT22 by size exclusion chromatography (SEC) and crosslinked the sample using the MS cleavable Disuccinimidyl Dibutyric Urea (DSBU) crosslinker (Figure 1B-C and Supplementary Figure S1). DSBU is an amine- and hydroxy-specific, homo-bifunctional crosslinker with a crosslinking spacer arm of 12.5Å (Iacobucci et al., 2018) that connects both closely packed residues within Cα-Cα Euclidean distances of 27Å and flexible residues located up to 43Å apart (Felker et al., 2021). A second protein complex comprising *Chlamydomonas* IFT-B2 proteins IFT57, IFT38, IFT54 and IFT20 as well as the IFT-B1 proteins IFT88 and IFT52N was also subjected to crosslinking/MS to provide data on the interactions within the IFT-B2 complex and between the IFT-B1 and B2 sub-complexes (Figure 1C). The crosslinking experiments of these two complexes were performed independently and were subsequently digested with both LysC and trypsin. The resulting peptides were enriched by strong cation-exchange chromatography and subjected to MS/MS analysis. Identification of crosslinking pairs was performed with the MeroX software (Götze et al., 2015) taking into account all possible crosslinks of DSBU. Only crosslinking data with false discovery rate (FDR) below 1% and scores above 80 are considered high-confidence and used in the analysis below.

Within the IFT-B1 nonamer, we identified 402 intra- and 859 intermolecular (Figure 1B, Supplementary material 1). Multiple intramolecular crosslinks were present within IFT88, IFT81, IFT74 and IFT70 (Figure 1B, brown lines). The intramolecular crosslinks of IFT88 show a 34-50 residues periodicity, which agrees with its predicted tetratricopeptide repeat (TPR) structure. The intramolecular crosslinking network of IFT81 and IFT74 shows a similar pattern with periodicities of 10-25, 50-80, 200-250 and 400-450 residues, suggesting that crosslinks formed within the same helix, and between adjacent coiled-coils (CCs). Together with intermolecular crosslinks between the N- and C-terminal halves of IFT81/74, these crosslinks are consistent with a heterodimeric IFT81/74 structure consisting of parallel CCs and agree with the crystal structure of *Trypanosoma* IFT81N/74N/22 previously published (Wachter et al., 2019). The IFT27/25 hetero-dimer primarily crosslinks to the C-terminal part of IFT81/74 while IFT22 crosslinks to the central part of IFT81/74 (Figure 1B). The C-termini of IFT46 and IFT52 interact in a hetero-dimer (IFT52C/46C) that was previously shown to mediate the interaction between IFT88/70/52/46 and IFT81/74/27/25/22 subcomplexes (Taschner et al., 2014). In our crosslinking data, IFT46C/52C crosslinks to the C-terminal part of IFT81/74 close to the IFT27/25 binding site. In addition, IFT81N/74N crosslinks to IFT88 and IFT70, which may constitute a second interaction site between IFT81/74 and IFT88/70/52/46 sub-complexes within the IFT-B1 complex. The most N-terminal 150 residues of IFT88 crosslink primarily to the 250 most C-terminal residues of IFT70 indicating an N- to C-interaction, while IFT52 crosslinks to both IFT70 and IFT88.

Analysis of the IFT-B1_B2 hexamer crosslinked sample revealed 575 intra- and 383 intermolecular high-confidence crosslinks (Figure 1C, Supplementary material 2). 78 intermolecular crosslinks were identified between IFT57 and IFT38 (Figure 1C) in agreement with previous data showing that these two proteins interact via their C-terminal CC domains (Taschner et al., 2016) (Figure 1A). This is also the case for IFT54 and IFT20 (Figure 1C) that form a complex via their C-terminal helices (Figure 1A). The N-terminal CH-domain of IFT54 is linked to the C-terminal CC domain via a long linker region that is presumably disordered and provides high flexibility in the relative position of these two domains of IFT54. This notion is supported by our crosslinking analysis where the CH-domain of IFT54 is crosslinked to the CCs of IFT54/20 along most of their lengths (Figure 1C). IFT57, like IFT54, contains a long intrinsically disordered central region between the N-terminal CH-domain and the C-terminal CCs. For IFT57, we also observe a crosslinking pattern where the IFT57 CH-domain form crosslinks to the C-terminal CCs of IFT57 and the binding partner IFT38 (Figure 1C). These results suggest a high degree of flexibility in the position of IFT54 and IFT57 CH-domains with respect to the CCs. Our MS analysis identified multiple crosslinking pairs formed between IFT88 and IFT52, IFT57 or IFT38 (25, 13 and 6 high-confidence crosslinks, respectively). These crosslinks suggest, in agreement with previously published results (Taschner et al., 2016), that IFT88/52 bridges IFT70 of the IFT-B1 complex to IFT57/38 of the IFT-B2 complex thus connecting B1 and B2 within IFT-B.

### Prediction and validation of the IFT81/74/27/25/22 structure

IFT81 and IFT74 associate into a hetero-dimer via CCs and serve as a scaffold onto which the small Rab like GTPases IFT22, IFT27 and RabL2 associate (Kanie et al., 2017; Nishijima et al., 2017; Taschner et al., 2014; Wachter et al., 2019). RabL2 only associates with the IFT trains during the initiation and early steps of anterograde IFT (Kanie et al., 2017) and was not included in the current study. Formation of the IFT81/74 hetero-dimer is essential for IFT in *C. elegans* (Kobayashi et al., 2007) and is a pre-requisite for IFT train assembly at the ciliary base (Brown et al., 2015). IFT25 is also loaded on the IFT81/74 platform via direct interaction with IFT27 (Bhogaraju et al., 2011). The N-termini of IFT81 and IFT74 were shown to bind tubulin heterodimers as cargo via a CH-domain and a positively charged region, respectively (Bhogaraju et al., 2013). The structure of the N-terminal half of *Trypanosoma brucei* IFT81/74 in complex with IFT22 was determined by X-ray crystallography and shows that IFT81N/74N organizes into 6 parallel CCs (CC I-CC VI), where IFT22 associates with CC VI (Wachter et al., 2019). Although the exact binding site is currently not known, IFT27/25 was shown to bind the C-terminal half of the IFT81/74 complex in both *Chlamydomonas* and human cells (Taschner et al., 2014; Zhou et al., 2022).

We made use of the recent advances in protein structure prediction by machine learning as implemented in AlphaFold (AF) (Jumper et al., 2021) using a local installation as well as a Colab notebook implementation (Mirdita et al., 2022) to model the structure of *Chlamydomonas* IFT-B sub-complexes, which allowed us to assemble a structural model for the 15-subunit IFT-B complex. All structural models of protein complexes were modelled using the AF multimer version optimized for the structure prediction of multimeric protein complexes (Evans et al., 2022). The quality of the resulting AF models was initially assessed using the predicted local distance difference test score (pLDDT), which constitute a per-residue score reporting on the confidence of the local structure prediction. Structural predictions with pLDDT>70 indicate confident parts of the model (colored blue in pLDDT figures) whereas low confidence structural segments with pLDDT<50 likely represent intrinsic disorder (colored orange in pLDDT figures) (Evans et al., 2022; Stevens and He, 2022). To evaluate the accuracy of the relative positions of protein subunits within multimeric structures, the predicted alignment error (PAE) plots were inspected to ensure that protein-protein interface residues have low error scores (for example see Figure 2D). Importantly, all protein-protein structure models are supported by observations from at least one biochemical or biophysical technique.

**Figure 2:**
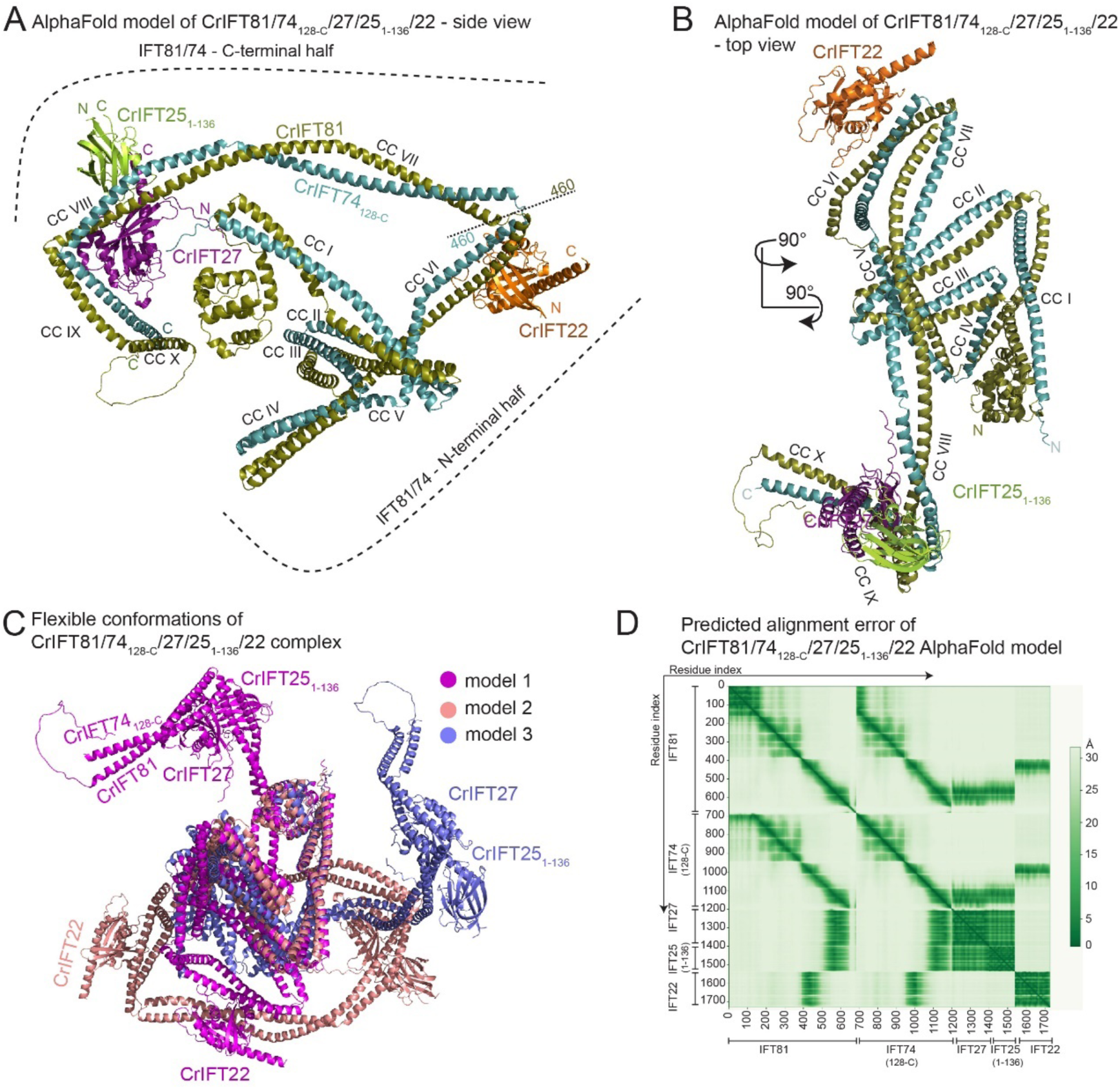
The AlphaFold predicted structure of CrIFT81/74/27/25_1-136_/22. **(A)** AlphaFold predicts the structure of IFT81/74_128-C_/27/25_1-136_/22 complex as two halves built on the IFT81/74 CC scaffold and hinged by a short loop region around the amino acids 460 of both IFT81 and IFT74. CrIFT22 binds near the hinge region on the CC domain VI (B) while the CrIFT27/25 heterodimer is docked proximal to the C-termini of IFT81/74 in a ‘L-shape’ structure formed by CCs VIII and IX. **(B)** The top view of the complex shows the organization of the IFT81/74 CCs and the docking sites for IFT27/25 and IFT22. **(C)** 3 independent AF models of the IFT81/74_128-C_/27/25_1-136_/22 complex are superimposed using their N-terminal halves to illustrate high flexibility between the separate half of IFT81/74 **(D)** The predicted alignment error (PAE) plot for the model shown in panels A-B. This plot assesses the confidence in the relative position of subunits within the complex. The Y- and the X-axis shows the residues indexed of the corresponding subunits as indicated. The aligned error in angstroms (Å) is color coded according to the bar to the right of the plot. Green color indicates low PAE (high confidence) and white color indicates high PAE (low confidence).

The structural model of the pentameric IFT81/74/27/25/22 complex shows that the IFT81/74 complex folds as 10 parallel CCs (CC I-CC X) connected by short loops (Figure 2A-B). Although CC VII-VIII and IX-X could be considered as single CC segments, resulting in a total of 8 CCs in IFT81/74, we denote these as separate CCs as we observe significant bends and/or breaks in the CC helices. Preceding the CCs are 130 residues of IFT81 that adopt the fold of a CH domain and 100 residues of IFT74 are predicted to be unstructured (Figure 2A-B) (Bhogaraju et al., 2013). The local structure of all 10 CCs is predicted with high confidence as highlighted by the coloring of the IFT81/74/27/25/22 model according to the pLDDT score (Figure S2A). Contrary, the structure of the short linker regions connecting adjacent CCs are predicted with lower confidence implying possible flexibility in the position between connecting CCs (Figure S2A). In particular, the loop region connecting CC V to CC VI has pLDDT<50 and may not adopt an ordered structure in solution. This hinge region divides the IFT81/74 complex into approximate N- and C-terminal halves (Figure 2A) and likely allows for a high degree of flexibility as no obvious protein-protein interface restricts the relative position of N- and C-terminal parts of IFT81/74. Indeed, although different structural models produced by AF have very similar local structures for the IFT81/74 N- and C-terminal parts separately, their relative position vary greatly (Figure 2C). This notion is also supported by the PAE plot that shows low errors for IFT81 and IFT74 helices of the same CC but much larger error between residues of N- and C-terminal CCs (Figure 2D). However, 269 crosslinks between residues of the N- and C-terminal halves of IFT81/74 suggest that the complex can adopt a compact conformation in solution where the two halves are in proximity (Figure 1B).

In our structural model, IFT22 and IFT27/25 are positioned on CC VI and CC VIII-CC IX of IFT81/74, respectively, with high confidence as illustrated by the low PAE for interacting regions (Figure 2D). The *Chlamydomonas* IFT81/74 model predicted here superimposes well onto the *Trypanosoma brucei* IFT81N/74N/22 crystal structure (Figure S2B). The binding site of IFT22 on CC VI of IFT81/74 thus appears to be conserved between *Chlamydomonas* and *Trypanosoma*. We further validated the structural model by site directed photo-crosslinking using a purified *Chlamydomonas* IFT81/74/27/25_1-136_/22 complex where the native amino acid E418 of IFT81 located near the IFT22 binding site was substituted with the UV-reactive amino acid p-benzoyl-L-phenylalanine (pBpa) (Figure S2C) (Young et al., 2010). Upon UV activation pBpa forms a covalent bond with proteins located in the immediate vicinity allowing the crosslinked proteins to be resolved on SDS-PAGE as they migrate slower than their monomeric non-crosslinked counterparts. The IFT81/74/27/25_1-136_/22 complex containing pBpa at position 418 in IFT81 formed a stoichiometric, UV dependent crosslink with IFT22 that migrated on SDS-PAGE at expected molecular weight of 123 kDa (Figure S2C). We tested the site-directed specificity of the method by using another IFT81/74/27/25_1-136_/22 complex that contained pBpa in the CH-domain of IFT81 (position 68), far away from the predicted IFT22 binding site. This complex did not form crosslinks with IFT22 upon UV activation. These data provide strong experimental evidence for the position of CrIFT22 on CC VI of CrIFT81/74 as illustrated in Figures 2 and S2.

IFT81_534-623_/74_533-615_ encompasses CCV III and CC IX and adopts an L-shaped structure that cradles the IFT27/25 heterodimer with IFT27 contacting both CC VIII and CC IX of IFT81/74 and IFT25 contacting only CC VIII (Figure 2A and S3). This binding site is consistent with the intermolecular crosslinking data obtained from the IFT-B1 complex (Figure 1B). A total of 41 crosslinks formed by IFT25 with IFT81/74 were found, of which 20 crosslinks were mapped to the N-terminal half and 21 to the C-terminal half of IFT81/74. All IFT25 crosslinks with the C-terminal half of IFT81/74 were mapped to the IFT81_534-623_/74_533-615_ region. IFT27 made 59 crosslinks with IFT81/74 of which 35 were identified within the N-terminal part and 24 within the C-terminal part of IFT81/74. 19 out of 24 crosslinks between IFT27 and the C-terminal half of IFT81/74 were mapped to the IFT81_534-623_/74_533-615_ region. For a 3D visualization of the crosslinking network, we labelled the IFT25 (Movie 1) and IFT27 (Movie 2) crosslinks onto the IFT81C/74C model. The fact that IFT27/25 also crosslinks with the N-terminal half of IFT81/74 suggest that the N- and C-terminal halves can be in proximity within the complex consistent with a high degree of conformational flexibility as noted above. Although IFT27/25 crosslinks to both N- and C-terminal halves of IFT81/74, the main binding site on CC VIII-IX was verified experimentally as IFT81/74 proteins lacking the N-terminal 459 residues still associate with IFT27/25 (Figure 4C-D). This notion is in agreement with previous biochemical studies showing that CrIFT27/25 does not bind to the N-terminal IFT81_133–442_/74_135–475_ complex (Taschner et al., 2014). Thus, we conclude that the main docking site of IFT27/25 is on the C-terminal half of IFT81/74 in agreement with the predicted structural model of the pentameric IFT81/74/27/25/22 complex (Figure 2A). The data show that IFT27/25 and IFT22 have unique binding sites on the IFT81/74 scaffold and suggest a high degree of conformational flexibility between the N- and C-terminal halves of the IFT81/74 complex.

### Structural model of the IFT88/70/52/46 IFT-B1 sub-complex

We previously showed that IFT52 functions as a central IFT-B protein that connects IFT88, IFT70 and IFT46 in a tetrameric IFT-B1 sub-complex (Taschner et al., 2011). Subsequent structural studies revealed that the TPRs of IFT70 wrap around a proline rich region of IFT52 (residues 330-370) that constitutes the hydrophobic core of IFT70 (Taschner et al., 2014). Proximal to the IFT70 binding site, IFT88 contacts residue 281-329 of IFT52 (Taschner et al., 2014). In pull-down assays, residues 118-437 of IFT88 were sufficient for IFT52 interaction (Taschner et al., 2014). In addition, human IFT70 was shown to interact with the IFT88/52 dimer by visual immunoprecipitation assays and this interaction is essential for ciliogenesis (Takei et al., 2018). It is thus firmly established that IFT88/70/52/46 form a tetrameric complex although high-resolution structures are only available for *Chlamydomonas* IFT70/52 and *Tetrahymena* IFT52/46 (Taschner et al., 2014) and it is currently unknown how IFT88 interacts with IFT70/52.

Using AF, we predicted the structure of *Chlamydomonas* IFT88/70/52/46 in complex with the very C-terminal helices (CCX) of IFT81/74 (Figure 3A). This structural model is predicted with high confidence, except for a few flexible loops and termini (Figure S4A), as evident from the high pLDDT score and the low PAE values for interacting residues of all protein-protein interfaces (Figure 3B). Residues 120-713 of IFT88 are predicted to fold into 15 TPRs with the most N-terminal 119 and the most C-terminal 67 residues predicted to be intrinsically disordered (Figure 3A and 4E). IFT88 adopts a rather loose and open superhelical structure, in contrast to the tight and closed superhelical structure of IFT70 that buries residues 330-370 of IFT52 (Figure 3A and 3D). The interaction of IFT88 with IFT52_1-329_ can be divided into two main interfaces. For the first interaction site, the three most C-terminal TPRs of IFT88 adopt an extended conformation to interact with the N-terminal GIFT domain of IFT52 (residues 1-270) with 40 predicted high-confidence (PAE<5Å) close contacts within 5Å. For the second interaction site, residues 271-330 of IFT52 interact in an extended conformation with the most N-terminal 12 TPRs of IFT88 (Figure 3A). The following 330-370 residues of IFT52 snake their way through the interior of the IFT70 superhelix as previously observed in the IFT70/52 crystal structure (Taschner et al., 2014). Finally, the small C-terminal domain of IFT52 (residues 371-C) interacts with the C-terminal domain of IFT46 to form a small hetero-dimer at the N-terminal face of the IFT70 superhelix (Figure 4E). In this structural model, the four most N-terminal TPRs of IFT88 are within interaction distance of the three most C-terminal TPRs of IFT70 (Figure 3C) in agreement with the direct IFT88-IFT70 interaction observed in pull-down experiments (Taschner et al., 2014). However, there appears to be no direct non-covalent interaction to tether IFT52C/46C to the N-terminal face of IFT70 (Figure 3A and 3E). We conclude that IFT52 is a central hub that organizes the IFT-B1 complex, which explains why *Chlamydomonas* ift52 mutant cells (bld1) contain highly destabilized IFT-B1 complexes (Richey and Qin, 2012) and display severe ciliogenesis defects (Brazelton et al., 2001).

**Figure 3:**
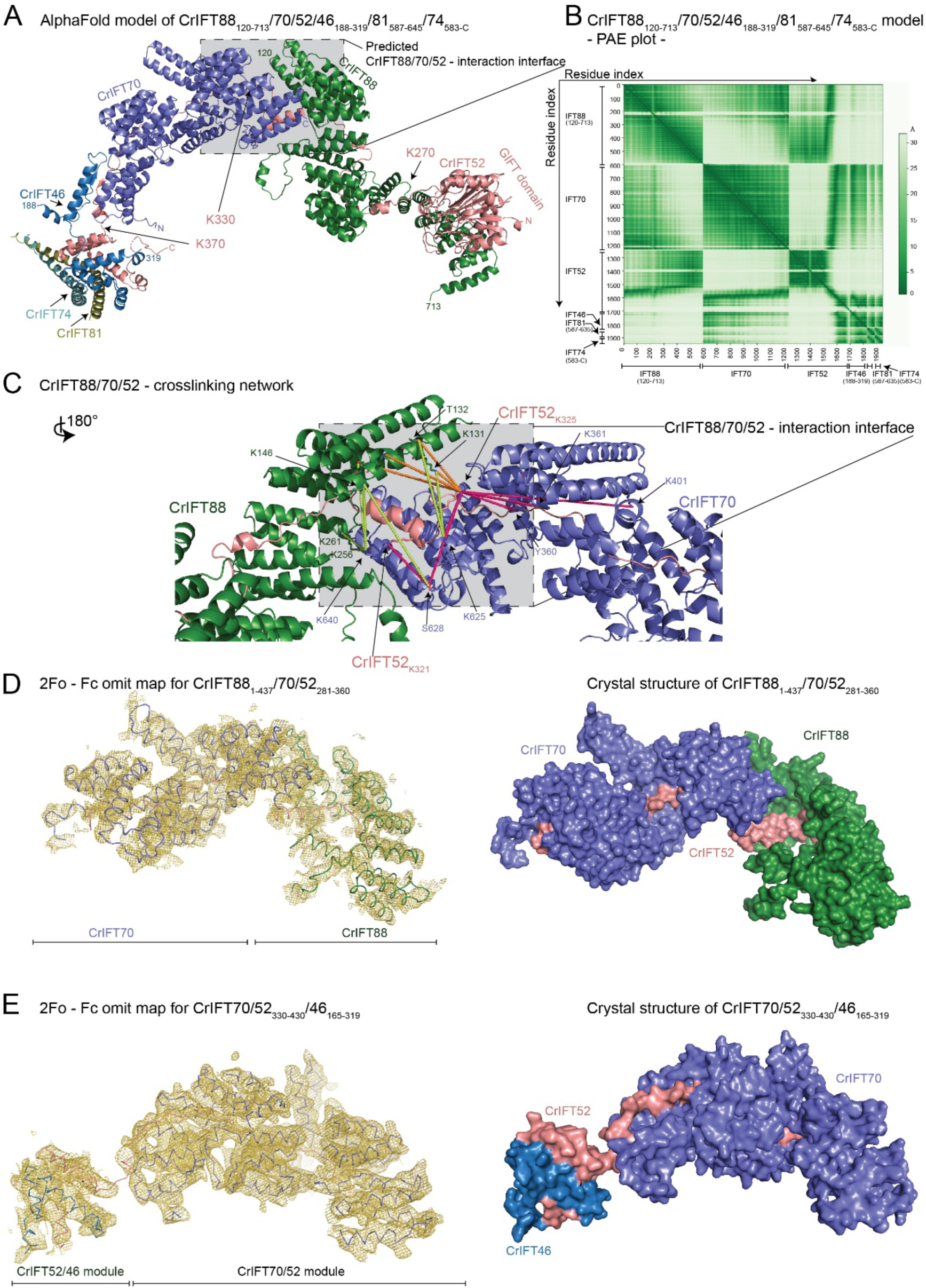
Structural model of the IFT88/70/52/46 IFT-B1 sub-complex. **(A)** The AlphaFold predicted model of CrIFT88_120-713_/70/52/46_188-319_/81_587-645_/74_583-C_. **(B)** The predicted alignment error plot of the complex from A. The residue indexes are indicated on the X- and Y-axis. **(C)** The CrIFT88/70/52 crosslinking network validates the interaction interface predicted by AlphaFold. The lime-green dashed lines are showing crosslinking pairs formed between IFT88 and IFT70. The orange dashed lines are showing the IFT88/52 crosslinks and the hot pink lines the crosslinks between IFT52 and IFT70. K321 and K325 of IFT52 make multiple short-range interactions with both IFT88 and IFT70 residues. **(D)** The crystal structure of *Chlamydomonas* IFT88_1-437_/70/52_281-360_ displayed as ribbon and the 2Fo - Fc omit map as a yellow mesh (left panel). Surface representation of the structure is shown on right panel. **(E)** The crystal structure of *Chlamydomonas* IFT70/52_330-430_/46_165-319_ displayed as ribbon and the 2Fo - Fc omit map as a yellow mesh (left panel). Surface representation of the structure is shown on right panel.

**Figure 4:**
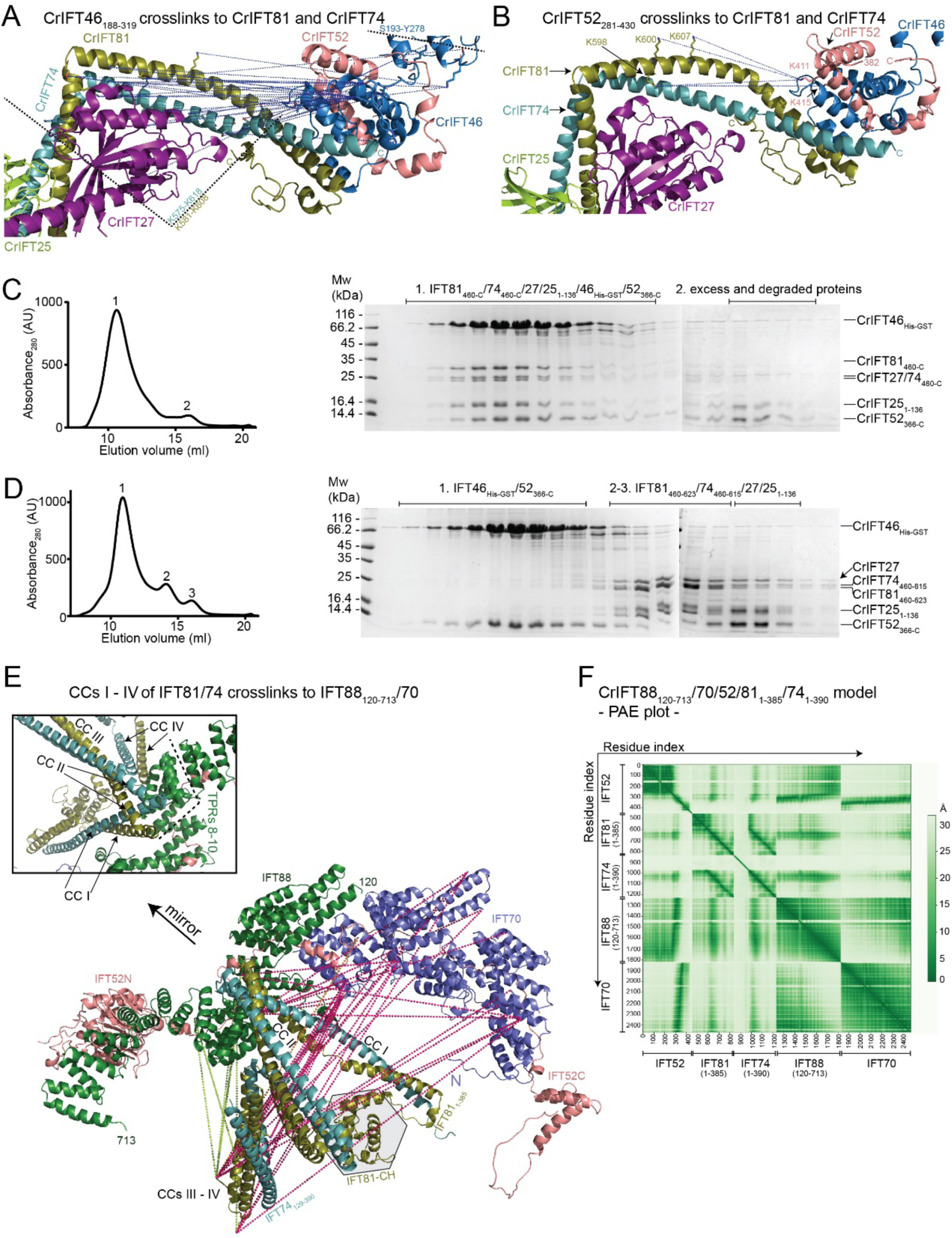
IFT81/74 has two separate binding sites on IFT88/70/52/46. **(A)** The blue dotted lines indicate the IFT46 crosslinks to the C-termini of IFT81 and IFT74. These crosslinks map to a 43 residue stretch from K575 to K618 in IFT81 and a 27 residue stretch in IFT74 bordered by K581 and K608. An additional crosslinking pair was identified between IFT46 and IFT27. **(B)** K411 and K415 of IFT52 crosslink to K598, K600 and K607 of IFT81. These crosslinks are labelled as in panel (A). **(C-D)** The C-terminal domains of IFT52 and IFT46 co-purify with the IFT81_460-C_/74_460-C_/27/25_1-136_ protein complex (C) but not with a protein complex that is missing residues 623-C of IFT81 and 615-C of IFT74 (D). The SEC fractions indicated on the right with 1-3 were analysed by SDS-PAGE and stained with Coomassie to evaluate the protein composition. **(E)** IFT81_1-385_/74_128-390_ crosslinking network with IFT88, IFT70 and IFT52 labelled on the AF predicted model of *Chlamydomonas* IFT88_120-713_/70/52/81_1-385_/74_1-390_. The N-terminal 128 amino acids of IFT74 are predicted to be unstructured and omitted from the figure for clarity. The box shows a zoom-in on the IFT81/74/88 interaction interface. **(F)** The predicted alignment error plot of the *Chlamydomonas* IFT88_120-713_/70/52/81_1-385_/74_1-390_ complex.

We validated the structural model of IFT88/70/52/46 using crosslinking/MS and crystallographic X-ray diffraction data. Several crosslinks are found at the IFT88/70/52 interaction interface. Lysines 321 and 325 of IFT52 crosslink to residues of both IFT70 and IFT88 in agreement with a composite interaction interface (Figure 3C and Movie 3). In addition, lysines 625 and serine 628 of IFT70 make multiple crosslinks to residues in the N-terminal part of IFT88 (residues 131-262, Figure 3C and Movie 3). To further validate the predicted structural model, crystals were obtained for a minimal IFT88_1-437_/70/52_281-360_ complex and X-ray diffraction data were collected to 3.8Å resolution (Supplementary Table 1). Molecular replacement with the IFT70/52 crystal structure (Taschner et al., 2014) and the AF model of the IFT88_120-437_ fragment gave a unique solution (Top LLG of 832) and the resulting omit electron density map clearly identifies the position of IFT70 and IFT88 TPRs and validates the position of the interacting regions of IFT88 and IFT70 within the complex (Figure 3D).

No crosslinks between the IFT52C/46C heterodimer and IFT70 were observed in our crosslinking/MS dataset. However, published crystal structures of CrIFT70/52 and *Tetrahymena thermophila* (Tt)IFT52C/46C (Taschner et al., 2014) and the fact that only 4 residues covalently link the part of IFT52 emerging from the IFT70 superhelix to the C-terminal domain of IFT52 that interacts with IFT46C effectively restrain the relative position of IFT70 and IFT52C/46C within the complex (Figure 3E). However, to validate the IFT70/52/46 structural model, we crystallized a minimal CrIFT70/52_330-430_/46_165-319_ complex and collected X-ray diffraction data to a resolution of 4Å. The crystal structure was determined by molecular replacement using the previously determined crystal structure of CrIFT70/52 (Taschner et al., 2014) and an AF generated model of *Chlamydomonas* IFT52C/46C, which resulted in a unique solution. The resulting omit electron density map clearly positions the IFT52C/46C complex at the N-terminal face of IFT70 (Figure 3E). However, given that IFT52C/46C is connected to IFT70 via a 4-residue covalent linker of IFT52 with no non-covalent interactions observed, the position of IFT52C/46C relative to IFT70 is likely quite flexible to accommodate different conformations in solution. In summary, a combination of AF modelling, chemical crosslinking and X-ray crystallography support the structural architecture of the IFT88/70/52/46 complex shown in Figure 3.

### IFT81/74 can associate with IFT88/70/52/46 via two distinct interaction sites to form the IFT-B1 complex

With validated structural models of IFT81/74/27/25/22 and IFT88/70/52/46 (Figures 2–3) in hand, we wanted to address how these two sub-complexes associate to form the IFT-B1 complex. We previously showed that the IFT52C/46C complex co-purified with IFT81ΔN/74ΔN/27/25 for both *C. reinhardtii* and *T. thermophila* and mapped the interaction to the C-terminal half of the IFT81/74 complex (Taschner et al., 2014). Recently, it was shown that human IFT52/46 associates with the C-terminal part of IFT81/74 (Zhou et al., 2022) demonstrating evolutionary conservation for this interaction. The exact binding site of IFT52C/46C on IFT81C/74C is unknown and no structural information is available for the complex. To this end, we utilized the modelled structures of the C-terminal part of IFT81/74 together with the C-terminal domains of IFT52 and IFT46 (IFT81_460-C_/74_460-C_/27/25_1-136_/46_148-328_/52_382-C_, see Figures 2E and S3) and the modelled structure of the IFT88/70/52/46 tetramer together with the C-terminal CCs of IFT81/74 (Figure 3A-B). Both complexes are modelled with high confidence as demonstrated by pLDDT and PAE plots (Figures 2D, 3B and S3) and reveal identical binding sites for IFT52C/46C on the most C-terminal CCs (CC X) of IFT81/74. The interaction interface is distal to the IFT27/25 binding site, is mostly hydrophobic in nature, and is formed by the residues 623-654 of IFT81, 615-641 of IFT74, 235-319 of IFT46 and 371-454 of IFT52 (Figure 4A-B, S9C). When mapping the crosslinking pairs of IFT46 (Figure 4A) or IFT52 (Figure 4B) onto the predicted IFT81/74/27/25 structure, we observe that IFT52C/46C predominantly crosslinks to the IFT81_581-608_/74_575-618_ region, which constitute the IFT27 binding site (Figures 4A-B). The crosslinking pairs are thus mostly formed proximal to the interaction site predicted by AF suggesting either the predicted model is inaccurate or is perhaps a high availability of free amine residues that can facilitate crosslinking formation. We addressed these possibilities experimentally in interaction studies of IFT52/46 and 81/74 complexes either with or without the predicted interacting helices of IFT81/74 (IFT81_460-C_/74_460-C_/27/25_1-136_ or IFT81_460-623_/74_460-615_/27/25_1-136_). The results show that the IFT81_623-C_/74_615-C_ helices predicted to interact with IFT52/46 are indeed required for complex formation on SEC (Figure 4C-D) thus validating the structural model shown in Figure 4A-B.

Interestingly, we observed numerous crosslinks between IFT81N/74N and the opposite end of the IFT-B1 complex, meaning IFT88, IFT70 and the N-terminal GIFT domain of IFT52 (Figures 1B, S8D). This observation suggests a possible second binding site through which IFT81N/74N link more closely to IFT52/88/70. Indeed, AF predictions where the N-terminal sequences of IFT81 and IFT74 were used as input together with IFT88/70/52 (IFT81_N-385_/74_N-390_/52/70/88_121-713_) provided a structural model suggesting a second conformation of the IFT-B1 complex. In this second IFT-B1 conformation, the first 4 CC domains of IFT81/74 form a tetrahedral structure that interacts directly with IFT88 (Figure 4E). Specifically, the tips between CCs I – II and CCs III - IV of IFT81/74 binds adjacent to the TPRs 8-10 of IFT88 (residues 468-536) to form a 54 pair high-confidence (PAE<5Å) interaction interface within 5Å apart (Figure 4F). Conservation analysis also corroborates this finding as the region of binding on IFT81/74 is conserved (Figure S9B). To address if the two observed binding modes can happen simultaneously or are mutually exclusive, we used AF and IFT-B1 protein sequences where both binding sites are present (IFT81/74/52_251-C_/88_120-C_/70/27/25/22/46_188-320_). We produced 5 AF models of this IFT-B1 complex as well as two additional quintuples that allow for more flexibility by splitting the IFT81/74 in N- and C-terminus halves either by introducing a break in the polypeptide chains or via a 100-residue glycine linker at position 458 in both IFT81 and IFT74. Of the 15 resulting models, 12 have the IFT81/74-C-terminus interaction with IFT46 and 7 models include both binding sites. Just one model identifies only the N-terminus IFT81/74 interaction with IFT88. However, none of five AF structures, where the native IFT81/74 sequences were used, had properly modeled α-helical structure of the central CCs but instead, at least some of the CC domains, resemble disordered regions. This suggest, that although no steric clashes prevent simultaneous binding of IFT81/74 to IFT88/70/52/46 via the two separate binding sites, it may require unfolding of CC segments and thus be unfavorable.

### IFT54/20 and IFT57/38 of the IFT-B2 complex form an anti-parallel hetero-tetramer

Previous studies suggest that IFT57, 54, 38 and 20 form a tetrameric complex composed of IFT54/20 and IFT57/38 hetero-dimers (Baker et al., 2003; Follit et al., 2006; Omori et al., 2008; Taschner et al., 2016). All four proteins contain C-terminal CC domains while IFT38, 54 and 57 also harbour N-terminal CH-domains. The only experimental structures currently available for the IFT57/38/54/20 complex is of the IFT54 CH-domain of both *Mus musculus* and *Chlamydomonas reinhardtii*. Both the IFT81 and IFT54 CH-domains, but not the IFT57 or IFT38 CH-domains, were shown to bind to αβ-tubulin *in vitro* (Bhogaraju et al., 2013; Taschner et al., 2016). The N-terminal CH-domains of IFT57 and IFT54 are connected to their C-terminal CC region by long and partly unstructured linkers (Taschner et al., 2012). Using AF, we predicted the structure of the CrIFT57/38/54_135-C_/20 tetramer (Figure 5A) and mapped crosslinking pairs onto the model (Figure 5C-D). The model is predicted with high confidence as revealed by pLDDT scores >90 (Figure S5) except for the long unstructured linkers that connect the CH- and CC-domains within IFT57 and IFT54 (these regions were removed in Figure 5C-D for clarity). PAE plots support the structural arrangement of the CC helices of IFT57/38/54/20 and furthermore suggest a well-defined position for the IFT38 CH-domain on the IFT54/20 two-bundle CC close to the tetrameric interface (Figure 5B), which is supported by 26 short-range (<32Å) crosslinking pairs (Figure 5D). The structural model reveals that the IFT57/38 and IFT54/20 hetero-dimers are formed by parallel helices of the CC domains (Figure 5A). The IFT57/38 and IFT54/20 hetero-dimers engage in an anti-parallel fashion so that the N-terminal ends of the four CC helices form a four-helix bundle (Figure 5C). Interestingly, the very N-terminal part of IFT20 is predicted to fold into a small 2-stranded anti-parallel β-sheet that packs against IFT57/38 to induce a bend in the CC helices and likely constitutes a hinge point for conformational flexibility (Figure 5C, S5). The structural arrangement of the parallel IFT57/38 and IFT54/20 hetero-dimers are supported by crosslinking pairs between numerous residues (Figure 5C-D). In addition, multiple crosslinks between all 4 subunits strongly support the anti-parallel assembly of the IFT57/38 and IFT54/20 hetero-dimers into a four-helix bundle. The association of the IFT38 CH-domain with IFT54/20 likely strengthens the tetrameric assembly. However, both the low pLDDT score and the high PAE score for the long linker regions connecting IFT54 and IFT57 CH-domains with their respective CC domains suggest that these are unstructured and likely provide a high degree of flexibility to the position of these CH domains within the IFT-B complex.

**Figure 5:**
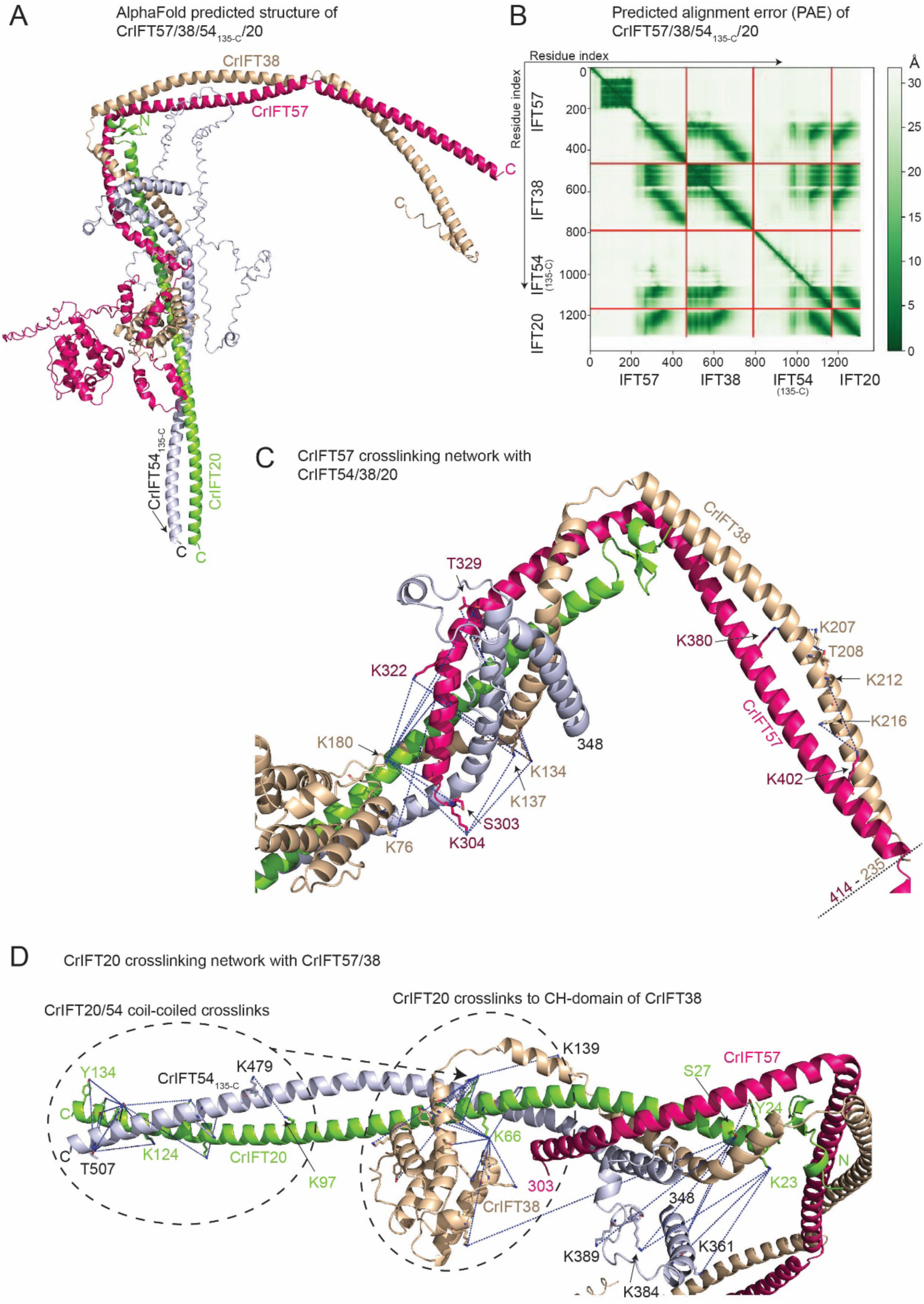
The AF predicted structure of the IFT57/38/54_135-C_/20 complex. **(A)** The AlphaFold predicted model of the *Chamydomonas* IFT57/38/54_135-C_/20 complex colored by chain. **(B)** Predicted alignment error of the AlphaFold model from (A). **(C)** Short-range crosslinking pairs formed by the helical domain of IFT57 with IFT54, IFT38 and IFT20 mapped as blue dotted lines between contributing sidechains. **(D)** Short-range crosslinking pairs of the IFT20 with IFT57, IFT54 and IFT38 labelled as blue dotted lines between the contributing side chains.

### IFT172 and IFT80 of the IFT-B2 complex interact directly

Previous studies have shown that IFT172 and IFT80 variants can result in skeletal ciliopathies and that these two subunits interact genetically (Boldt et al., 2016; Halbritter et al., 2013). However, IFT172 and IFT80 did not interact physically in sucrose gradients (Lucker et al., 2005) nor did they co-purify during SEC, which suggest that any direct physical interaction is relatively weak (Taschner et al., 2016). However, given that both IFT172 and IFT80 associate with the IFT57/38 heterodimer (Taschner et al., 2016), they could be in proximity within the IFT-B2 complex. To test this hypothesis, we used AF with full length sequences of CrIFT172 and CrIFT80, which reliably predicted the structure of the heterodimeric IFT172/80 complex (Figure 6A). The structural model of the IFT172/80 complex was predicted with high confidence (Figure S6A-B) within the respective interacting regions of the proteins and show that the TPR repeats within the residues 626-785 of IFT172 interact with the TPR repeats of IFT80 within residues 583-627 (Figure 6A). Under a conservative 5Å PAE threshold of the AF-predicted IFT80-IFT172 structure, we observed 26 residue pairs at a distance between 2.5-5Å and 180 pairs within less than 10Å apart. Residues 786-C of IFT172 are predicted to fold into TPRs but as they do not appear to participate directly in the interaction with IFT80 and are predicted with lower confidence (data not shown), they were omitted from the model shown in Figure 6A. Curiously, we did not observe homo-dimer formation of IFT80 using AF, which contrasts previous crystallographic analysis (Taschner et al., 2018). To assess the IFT172/80 complex formation experimentally, we monitored the interaction in pulldown assays. Purified Venus-tagged CrIFT80 was used to pull down purified full-length and a N-terminal deletion of CrIFT172 that lacks the predicted IFT80 interaction domains. The results show that CrIFT80 pulls down full-length IFT172 but not a C-terminal construct that lacks the interacting region predicted by AF (Figure 6B). Venus-tagged IFT80 clearly pulls down sub-stoichiometric amounts of IFT172 suggesting a weak interaction, which agrees with observation that the two proteins do not associate during SEC (Taschner et al., 2016).

**Figure 6:**
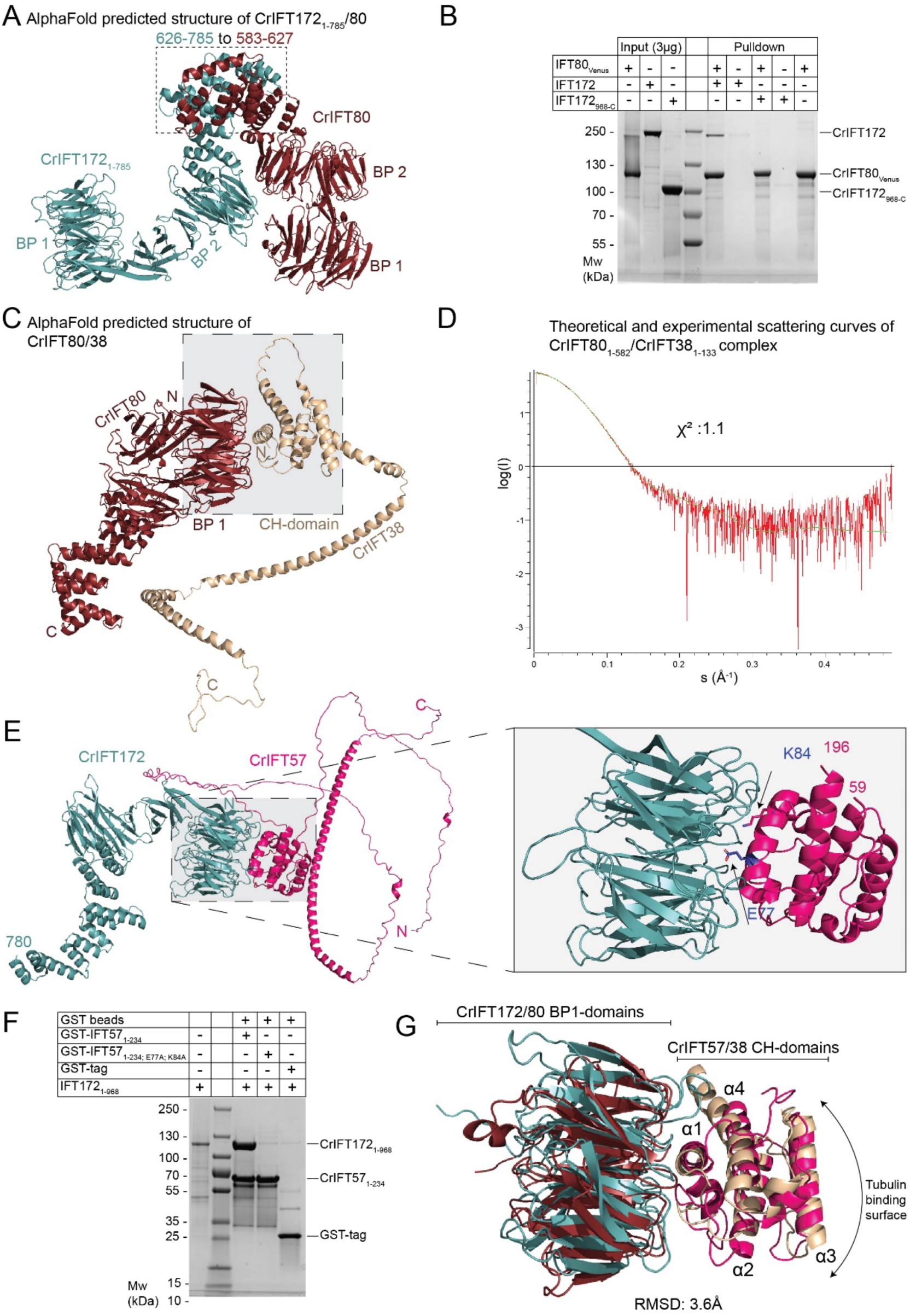
Dissecting the interaction between IFT80, IFT172, IT57 and IFT38. **(A)** AphaFold predicted structure of IFT172_1-785_/80. **(B)** Pull-down experiments with purified IFT80_Venus_ immobilized on GFP-binder beads and either IFT172 or IFT172_968-C_. IFT80 pulls down full length IFT172 but not the truncated version lacking the N-terminal 967 residues. **(C)** AlphaFold predicted structure of IFT80 in complex with IFT38 maps the interaction at the interface between the BP1 of IFT80 and the CH-domain of IFT38. **(D)** Comparison of the solution scattering curve of CrIFT80/38 as measured by SAXS and the calculated scattering curve for the IFT80/38 structural prediction. A ꭓ2 value of close to 1 indicate an excellent fit between measurement and calculation. **(E)** AlphaFold predicted structure of CrIFT172 and CrIFT57 shows interaction via the BP1 and CH domains. On the right panel, two residues in the CH-domain of IFT57, which are located at the interface with IFT172, are highlighted. **(F)** GST tagged CrIFT172_1-968_ pulls down the CH-domain of CrIFT57_1-234_ but not the mutated CH-domain of CrIFT57 where E77 and K84 residues were replaced by alanines. **(G)** Superimposition of the structure of BP1 of CrIFT80 (colored in red ruby) in complex with the CH-domain of 38 (colored in beige) with the BP1/CH domains of CrIFT172/57 (colored in teal and hot pink, respectively) shows a conserved mechanism of interaction different from the canonical tubulin binding mode exhibited by CH domains.

### IFT80 and IFT172 associate with the IFT-B2 complex through conserved β-propeller/CH-domain interactions with IFT38 and IFT57

Within the IFT-B2 complex, IFT80 and IFT172 were shown to associate with the CH-domains of IFT38 and IFT57, respectively (Taschner et al., 2018, 2016). CH-domains typically associate with microtubules/tubulin and/or actin along with a few other proteins involved in cellular signalling (Yin et al., 2020) and, to the best of our knowledge there is no structural characterization of how CH-domains associate with β-propellers. We thus used AF to model the structure of the respective interacting domains in CrIFT172/57 and CrIFT80/38 and validated the resulting models by small angle X-ray scattering (SAXS) and structure-guided mutagenesis.

The structural model of CrIFT80/38 shows that the N-terminal BP1 of IFT80 interacts with the CH-domain of IFT38 (Figure 6C). This interaction is mainly mediated by alpha helix α1 of the IFT38-CH domain that runs across the N-terminal face of the first β-propeller domain of IFT80 (Figure 6C, 6G). The IFT80/38 complex structure is predicted with high confidence as shown by the pLDDT scores >90 (Figure S6C) and the low PAE scores for interacting domains (Figure S6D). On the contrary, the C-terminal CC helix of IFT38 is folded with low confidence (pLDDT score <50), which likely reflects the absence of the interacting partner IFT57 in this model. To verify the validity of the model, we purified the IFT80_1-582_/38_1-133_ complex and collected SAXS data. The comparison of the theoretical X-ray scattering curve of IFT80_1-582_/38_1-133_ structural model with the experimental curve shows an almost perfect fit with a ꭓ2 value of 1.1 (Figure 6D) thus validating the structural model.

As was observed in the CrIFT80/38 structural model, AF predictions of the CrIFT172_1-780_/57 complex structure revealed an interaction between the first β-propeller of IFT172 and the CH-domain of IFT57 with high pLDDT and low PAE scores (Figures 6E and S6E-F). The interaction of IFT172 with the IFT-B complex was previously shown to be salt labile (Taschner et al., 2016), which agrees with the highly hydrophilic interface observed between IFT172 and IFT57 in our structural model. We used this structural model for mutagenesis designed to disrupt the IFT172/57 interaction interface (Figure 6F). The E77 and K84 residues of the CrIFT57 CH-domain lie in the interface with IFT172 BP1 and were mutated to alanine and used in pull-down experiments with purified CrIFT172_1-968_. The results show that while IFT172_1-968_ is pulled-down in stoichiometric amounts by wildtype CrIFT57_1-234_, the interaction is completely lost in the E77A, K84A double point mutant (Figure 6F). Taken together, these data indicates that IFT172 associates with the IFT-B2 complex through a strong interaction with the CH-domain of IFT57 and a rather weak interaction with IFT80.

Interestingly, the interaction involving the first β-propeller (BP1) of IFT172 and the CH-domain of IFT57 shows striking similarity to the IFT80/38 complex. IFT172/57 and IFT80/38 BP1-CH structural models superimpose with a root mean square distance (RMSD) of 3.6Å demonstrating a conserved interaction mechanism (Figure 6G). In both complexes, the interaction is mediated by charged residues of helix α1, which is markedly different from the tubulin binding mode exhibited by many CH-domains, which mainly involves residues in the vicinity of the corresponding helix α3 of the CH-domain (Bhogaraju et al., 2013; Hayashi and Ikura, 2003; Taschner et al., 2016) (Figure 6G). This likely points to β-propellers as a new class of interaction partner for CH domains in addition to the well-characterized tubulin and actin interaction partners.

### The IFT-B1-B2 connection

Previous biochemical data have shown that IFT88 and the N-terminal domain of IFT52 (residues 1-335) of IFT-B1 are sufficient to pull down the IFT-B2 complex via direct contacts to IFT57/38 (Katoh et al., 2016; Taschner et al., 2016). To gain structural insights into the IFT-B1-B2 connection, we used AF to model the structure of the IFT88/52N/57C/38C complex (Figure 7A). The structure of IFT88_120-713_/52_1-336_/57_356-C_/38_174-303_ was modelled with high confidence as revealed by high pLDDT scores for most of the model (Figure S7). The PAE plot (Figure 7B) demonstrates high confidence in the relative positions of all 4 proteins within the IFT88/52/57/38 complex. In the structural model, the C-terminal CC region of IFT57/38 engages IFT88/52, which creates a slightly arched structure where IFT88/57/38 loosely cradles the N-terminal GIFT domain of IFT52 (Figure 7A). The position of the IFT52 GIFT domain is supported by 11 crosslinking pairs with IFT88 and IFT57/38 as identified by MS (Figure 7C-D). Consistent with the structural model of IFT88/52/57/38 shown in figure 7A, two point mutations in the GIFT domain of IFT52 (K130E and R204E) that were previously published to significantly reduce the IFTB1-B2 interaction (Taschner et al., 2016) lie at the interface with IFT57/38 (Figure 7A). We also observe multiple crosslinks from IFT88_120-713_ to IFT38_212-C_ but no crosslinks to the IFT57_414-C_ region. Taken together, the crosslinking data supports the predicted structural model of CrIFT88_120-713_/52_1-336_/57_356-C_/38_174-C_ and reveals how IFT88/52 of IFT-B1 connect to IFT57/38 of IFT-B2.

**Figure 7:**
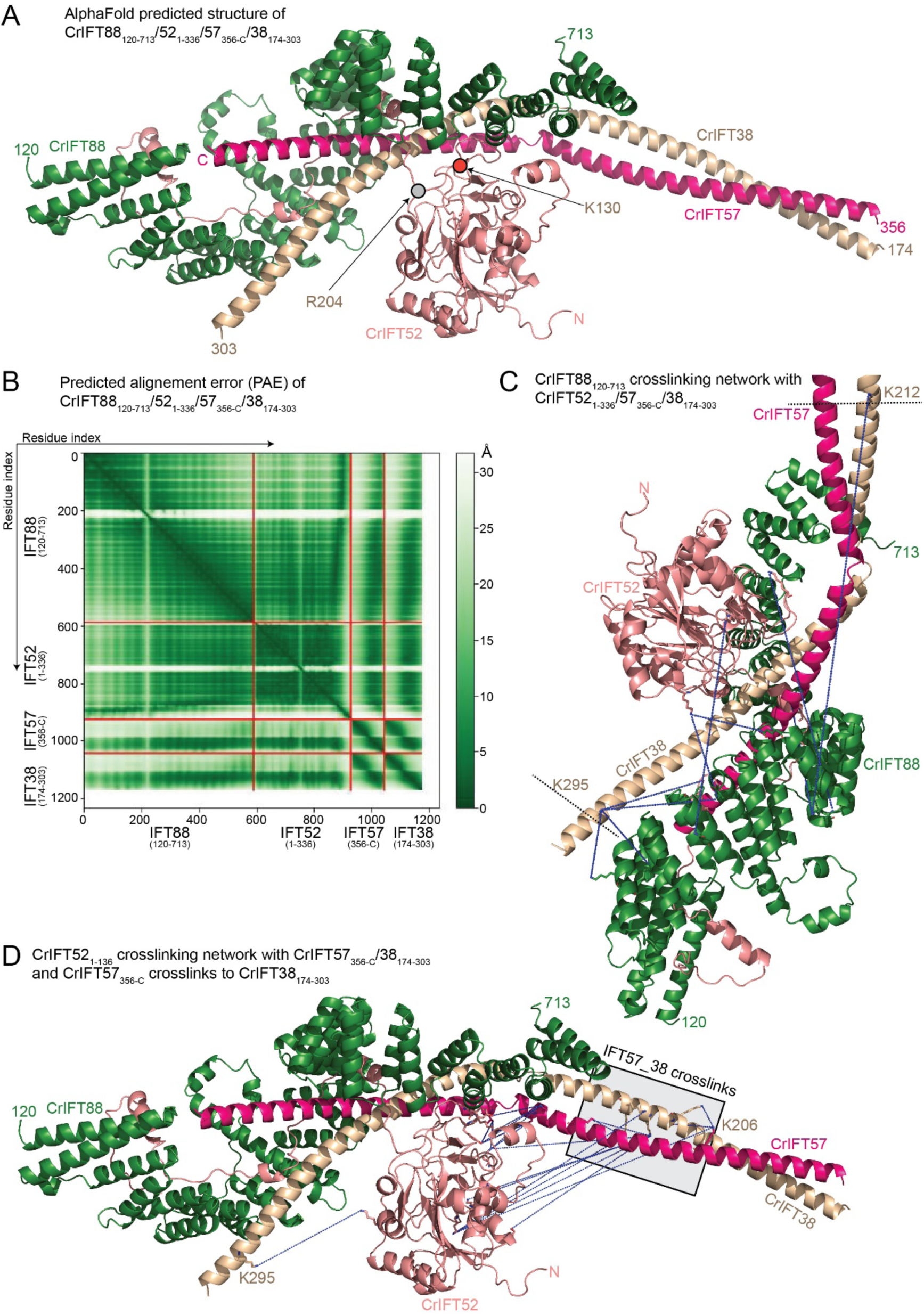
IFT88 links the IFT-B1 and IFT-B2 complexes via interactions with both IFT70 and IFT57/38. **(A)** AlphaFold predicted structure of the CrIFT88_120-713_/52_1-336_/57_356-C_/38_174-303_ complex. **(B)** Predicted alignment error (PAE) for the complex depicted in (A). **(C)** CrIFT88_120-713_ crosslinking network with CrIFT52_1-336_, CrIFT57_356-C_ and CrIFT38_174-303_ labelled with blue dotted lines. **(D)** The crosslinking interaction network of CrIFT52_1-336_ with CrIFT57_356-C_/38_174-303_ and of CrIFT57_356-C_ with CrIFT38_174-303_ labelled as in (C).

### Structural model of the 15-subunit *Chlamydomonas* IFT-B complex

From the experimentally validated structures of IFT-B sub-complexes displayed in Figures 2–7, we assembled the structural model of the 15 subunit *Chlamydomonas* IFT-B complex *in silico* (see Figure 8 and M&M). The structural model contains the proteins IFT81/74/27/25/22/52/46/88/70/57/38/54/20/80/172 but lacks RabL2 and IFT56. RabL2 is not a core member of the IFT complex as it dissociates shortly after departure of the anterograde IFT train from the ciliary base (Kanie et al., 2017; Nishijima et al., 2017). IFT56 is important for recruiting motility complexes to cilia and associates with the IFT-B complex via IFT46 and possibly IFT70 but does not appear to be essential to ciliogenesis in mice (Ishikawa et al., 2014; Swiderski et al., 2014). AF modeling places IFT56 close to IFT46 and IFT70 but as we were unable to express IFT56 as a soluble protein, we could not experimentally verify its position within the IFT-B complex and have thus omitted this subunit from our structural modeling. The model reveals an elongated IFT-B complex with a longest dimension of 430Å and a shortest dimension of 60Å (Figure 9A). These measures are consistent with the elongated IFT-B complexes organized into linear polymers with a repeat distance of 60Å that were observed observed in cryo-ET reconstructions of anterograde IFT trains (Jordan et al., 2018; van den Hoek et al., 2022).

**Figure 8:**
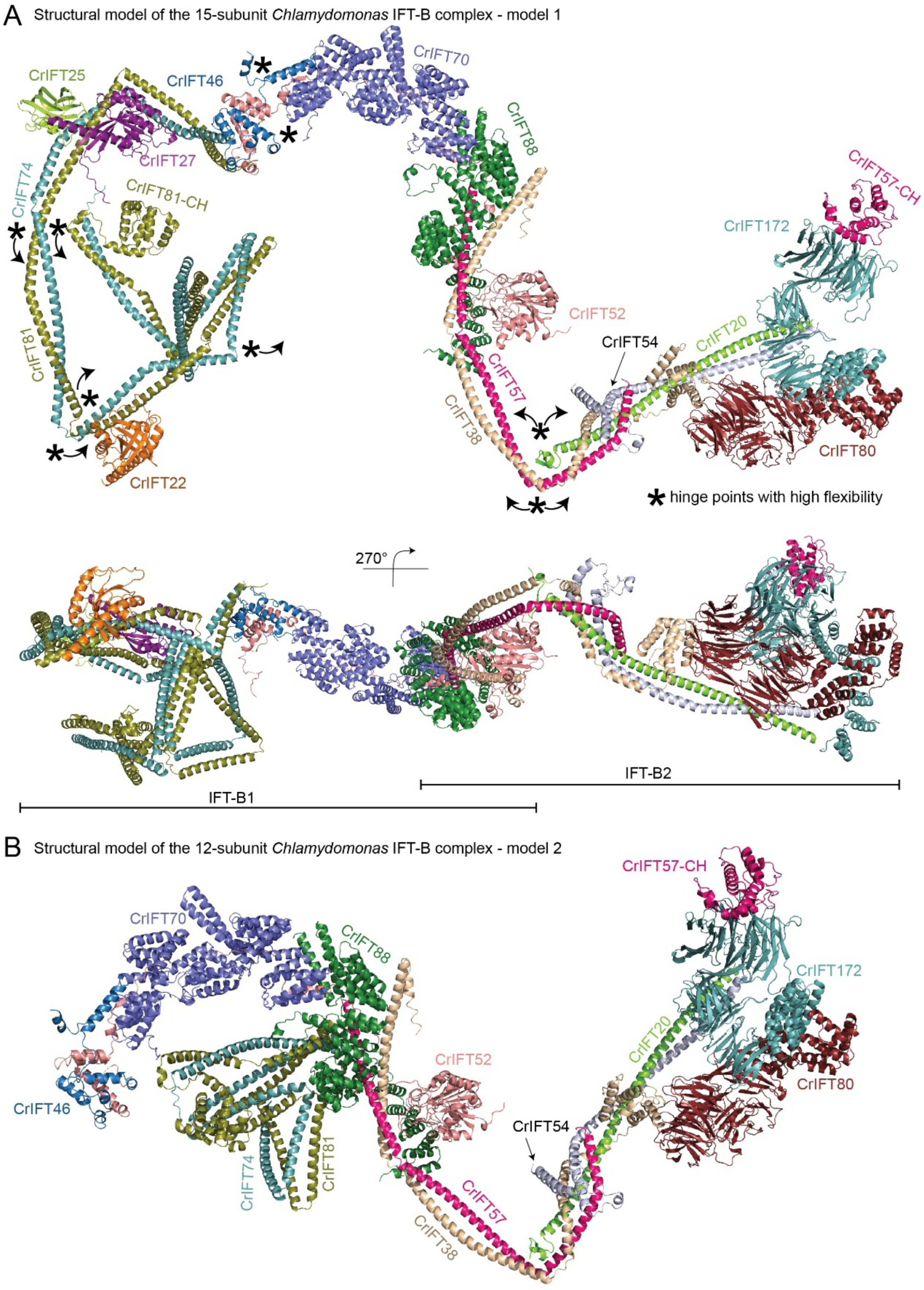
*In silico* structure of IFT-B 15-mer. **(A)** The model of 15-subunit *Chlamydomonas* IFT-B complex (model 1) assembled *in silico as* a rigid body from AF predicted sub-complexes using the binding site of IFT81/74 on IFT52/46 shown in Figures 3A and 4A-B. The structural model is assembled from CrIFT81/74_128-C_/27/25/22, CrIFT88_120-713_/70/52/46_188-319_/81_587-645_/74_583-C_, CrIFT88_120-713_/52_1-336_/57_360-C_/38_174-303_, CrIFT57/54_135-C_/38/20, CrIFT80/38 and CrIFT172_1-780_/57. **(B)** The model of 12-subunit *Chlamydomonas* IFT-B complex (model 2) assembled *in silico* using the IFT81/74 binding site on IFT88/70 shown in Figure 4E. The rigid body assembly was carried out using AF predicted structures of sub-complexes of CrIFT88_120-713_/70/52/46_188-319_/81_587-645_/74_583-C_, CrIFT88_120-713_/70/52/81_1-385_/74_1-390,_ CrIFT88_120-713_/52_1-336_/57_360-C_/38_174-303_, CrIFT57/54_135-C_/38/20, CrIFT80/38 and CrIFT172_1-780_/57.

**Figure 9:**
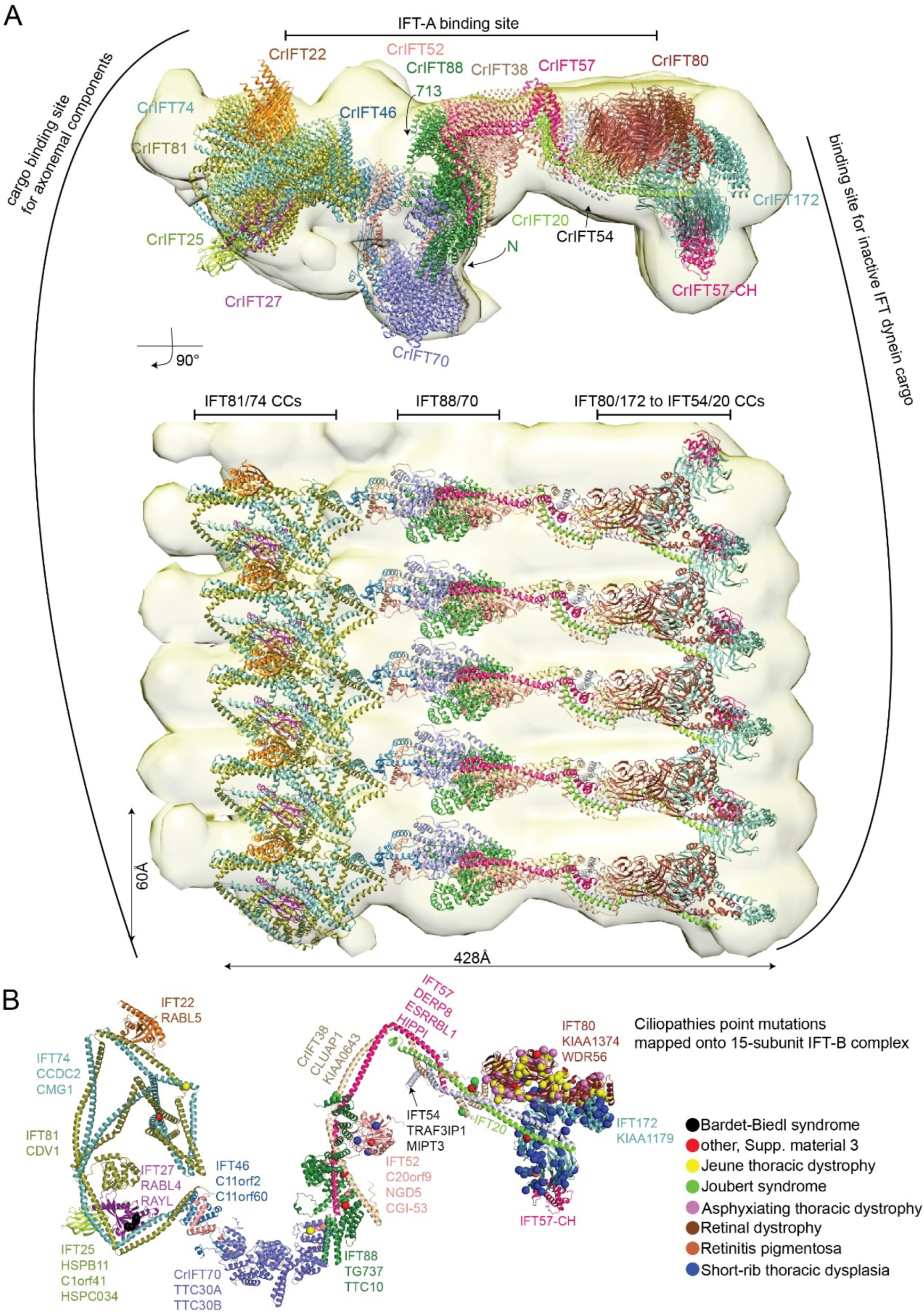
The IFT-B complex in context of anterograde IFT-B trains. **(A)** Molecular dynamic flexible fitting of the IFT-B 15-mer into the 25Å cryo-electron tomography map of the *Chlamydomonas* anterograde IFT-B trains obtained *in situ* (van den Hoek et al., 2022). The fit is shown in two perpendicular orientations. The highly elongated IFT-B complex fits with a repeat distance of 60Å consistent with anterograde IFT trains. **(B)** Single point mutations associated with ciliopathies are mapped onto the CrIFT-B 15-mer structure as spheres. Noteworthy, mutations that leads to amino acid deletions or additions, as well as frameshifts are not included in the figure. The IFT-B proteins are annotated according to their corresponding human gene names.

Several potential hinge regions are apparent in the structural model and likely serve as points of conformational flexibility within the IFT-B complex (marked with * in Figure 8A). Hinges are located between IFT70 and IFT52C/46C as well as between CC segments of IFT81/74 and IFT57/38 (Figure 8). To glean insights into conformational flexibility of the IFT-B complex in solution from the crosslinking/MS data, we labelled all intra- and intermolecular crosslinking pairs (Figure S8A-D). We considered crosslinking pairs with a distance <32Å as an indicator of structural rigidity and crosslinking pairs with distance >32Å as an indicator of conformational flexibility. The intramolecular crosslinking pairs suggest a high degree of conformational flexibility between the N- and C-terminal halves of the IFT81/74 complex (Figure S8A-B) consistent with our structural modeling (Figure 2C). Similarly, many intramolecular crosslinks with distances beyond 32Å were found between the N- and C-terminal ends of IFT38 and IFT57 (Figure S8B), which suggests a high degree of flexibility around the hinge point of the IFT57/38 CCs as indicated in Figure 8A. The intermolecular crosslinks show a similar pattern where numerous crosslinks between the two ends of the IFT57/38-54/20 hetero-tetramer suggest a significant bendability around the hinge region (Figure S8C-D). Intermolecular crosslinks are also very pronounced between IFT81N/74N and IFT88/70 and likely capture the second binding site for IFT81/74 on the IFT88/70/52/46 tetramer as highlighted in Figure 4E. The high degree of structural flexibility of the IFT-B complex is likely important for the polymerization into IFT trains and furthermore may provide a mechanism for the large structural rearrangements that occur when anterograde IFT trains remodel into retrograde IFT trains (Jordan et al., 2018).

Given the significant flexibility of the IFT-B complex in solution and the fact that the structural model of the 15 subunit IFT-B complex presented here was assembled by superposing smaller sub-complexes without the context of the IFT train, it is not surprising that a rigid-body manually fit the IFT-B 15mer into the cryo-ET map of anterograde IFT trains (van den Hoek et al., 2022) resulted in several subunits located outside of the density (data not shown). To obtain a better fit, we made use of the program Namdinator (Kidmose et al., 2019), which is an automatic molecular dynamic flexible fitting algorithm that requires only a structural model and a map as input. Two consecutive rounds of each 400.000 iterations in Namdinator resulted in a relatively good fit of the structural model of the IFT-B complex to the density of the anterograde IFT trains except for IFT22 and IFT25, which partly sit outside density (Figure 9A). One end of the elongated IFT-B complex contains the IFT-B1 complex with previously characterized cargo-binding sites for tubulin as well as outer-and inner dynein arms (Bhogaraju et al., 2013; Hou et al., 2007; Kubo et al., 2016; Taschner et al., 2017; Wang et al., 2020). The other end of the IFT-B complex harbors the IFT-B2 complex with IFT172, IFT80, IFT57CH and the C-terminal part of the IFT54/20 CCs positioned close to the binding site for the IFT dynein cargo on anterograde trains (Figure 9A). In the model, the IFT-A complex is positioned close to the CCs of IFT81/74, CCs of IFT57/38 and the C-terminal end of IFT88 (Figure 9A). However, due to the low resolution of the cryo-ET map (25Å) and the absence of the C-terminal domain of IFT172 in our model it is not possible to accurately pinpoint the IFT-B proteins involved in the interaction between IFT-A and IFT-B.

Our structural modeling and crosslinking/MS data revealed two separate binding sites of IFT81/74 on the IFT88/70/52/46 (Figure 4). AF was unable to model both sites simultaneously without disrupting and unfolding the CCs connecting the two halves of IFT81/74 and it thus appears likely that the two binding modes are mutually exclusive (for models of the IFT-B complex containing the alternative IFT81/74 binding mode see Figure 8B). Both models of the IFT-B complexes produce a relatively good fit to the 25Å cryo-ET map of the anterograde IFT train (using flexible fitting in Namdinator) and it is thus not clear from our data if one of these IFT-B models better represent the anterograde IFT train conformation.

## Discussion

### The *Chlamydomonas* IFT-B complex in context of IFT trains

We present an experimentally verified structural model of the 15 subunit IFT-B complex that is consistent with low-resolution cryo-ET reconstructions of anterograde IFT trains (Jordan et al., 2018; van den Hoek et al., 2022). However, given that our experimental validations were carried out on isolated IFT-B sub-complexes not polymerised into IFT-trains, our IFT-B structural model could represent a hybrid conformational state capturing conformations of both anterograde and retrograde IFT trains. During the preparation of this manuscript, three preprints using AF and cryo-EM to elucidate the structures of IFT-A and IFT-B complexes were published (Hesketh et al., 2022; Lacey et al., 2022; McCafferty et al., 2022). Lacey and co-workers elucidated the structure of anterograde IFT-trains at 10-18Å resolution and fitted AF generated models of IFT-A and B complexes into the density to obtain a pseudo-atomic model for the entire train structure (Lacey et al., 2022). Overall, the architecture of the IFT-B complex presented here agrees well with the cryo-ET structure presented by Lacey et al. Interestingly, in the anterograde IFT train structure, it is observed that IFT74N/81N associates with IFT88/70 (Lacey et al., 2022) consistent with the second IFT-B model described here (Figure 8B). In the cryo-ET structure, no density is observed for the C-terminal half of IFT81/74 associating with IFT27/25 and IFT22 suggesting that this part of IFT-B adopt flexible conformations and is likely averaged out in the cryo-EM maps (Lacey et al., 2022). The interaction between the C-terminal CC X domain of IFT81/74 and IFT52C/46C shown in Figure 4A-B are not observed in the anterograde IFT-B Cryo-EM structure.

If not important to the formation of anterograde IFT trains, what is the functional implication of the IFT81C/74C-IFT52C/46C interaction highlighted in Figures 4A-B and 8A? Firstly, IFT81C/74C-IFT52C/46C is a high affinity interaction (Taschner et al., 2014), which occurs through a conserved hydrophobic interface (Figure S9C). Indeed, the IFT81C/74C-IFT52C/46C interaction is evolutionarily conserved and was experimentally observed in *Chlamydomonas, Tetrahymena* (Taschner et al., 2014) and human (Katoh et al., 2016). Interestingly, our structural modelling by AF showed that significant unfolding of the IFT81/74 CC segments must occur for both IFT81/74 binding sites on IFT88/70/52/46 to be occupied simultaneously. This observation could suggest that the two binding modes are mutually exclusive and may happen separately in different cellular contexts. It is tempting to speculate that the IFT81N/74N-IFT88/70 structure shown in Figures 4E and 8A and the IFT81C/74C-IFT52C/46C structure shown in Figures 4A-B and 8A represent conformations unique to anterograde and retrograde IFT trains, respectively, although this notion can currently not be verified in the absence of cryo-ET reconstructions of retrograde IFT trains.

### Association of IFT-B with IFT-A and IFT motors

The IFT-B complex and its linear polymerization form the backbone of IFT trains onto which IFT-A polymers, dynein-1b, and finally kinesin-2 attach before entering the cilium (van den Hoek et al., 2022). How is IFT complex polymerisation into trains and association with motors facilitated? The fitting of the 15-subunit IFT-B structural model into a 5-repeat anterograde IFT-B train revealed that IFT-B polymerizes laterally and contains at one end the cargo binding sites for axonemal components and at the other end the binding site for inactive IFT dynein 1b cargo (Figure 9A). The lateral polymerization into trains appears involves 3 contact points provided by adjacent IFT81/74 CCs at one end, IFT88/70 complexes in the middle and the N-terminal CCs of IFT54/20, the N-terminal part of IFT172 and IFT80 at the other end (Figure 9A). This arrangement agrees with the recent anterograde IFT structure (Lacey et al., 2022). However, due to resolution limitations of IFT-A and IFT-B cryo-ET reconstructions that we used in this study, it is yet to be determined which residues within the 3-point-junction are essential for lateral polymerization of anterograde IFT trains.

Previous cryo-EM data have revealed a mismatch between the number of IFT-B, IFT-A and dynein 1b cargo complexes in anterograde IFT trains with approximate 6, 11.5 and 18nm repeat distances within the trains (Jordan et al., 2018; Toropova et al., 2019). IFT-A is flexibly tethered to IFT-B complexes through interactions between IFT139 of IFT-A and two copies of IFT81/74 of IFT-B at one end, IFT144/139 of IFT-A and the C-terminal TPR domain of IFT172 at the other end (Lacey et al., 2022). A third IFT-A/IFT-B interaction interface was elucidated by Hesketh and co-workers, who showed that the highly flexible IFT88 C-terminal extension bridges across to interact with IFT44 (Hesketh et al., 2022). Although we did not include the C-terminal part of IFT172 in our structural model of IFT-B, both IFT81/74 and IFT88 are positioned so that interactions to IFT-A proteins are favourable (Figure 9A).

Flexible fitting of the IFT-B model into cryo-ET maps of anterograde IFT trains suggest that the 60Å wide IFT-B complexes cannot support the loading of bulky IFT dynein cargo (Figure 9A) in agreement with previous published results (Jordan et al., 2018; Toropova et al., 2019). Lacey and co-workers showed that dynein cargo is using a composite surface formed by two adjacent IFT-B2 complexes. In our IFT-B model, IFT172, IFT80 and the CH domain of IFT57 supported by a shaft formed by the CCs of IFT54 and −20 are the main contributors for creating this composite binding site for dynein 1b cargo (Figure 8A and Figure 9A). Interestingly, the platform has a prominent negatively charged groove flanked by two positively charged regions formed by a conserved surface on BP1 of IFT172 and BP2 of IFT80 (Figure S9A-B).

Upon polymerization of IFT-A and IFT-B complexes at the base of cilia, the kinesin 2 motor associates with IFT-B to drive the anterograde IFT (van den Hoek et al., 2022). Because of its slim and flexible architecture, kinesin 2 is averaged out in cryo-ET reconstructions (Jordan et al., 2018; van den Hoek et al., 2022). However, biochemical studies showed that IFT88/57/52/38 (Funabashi et al., 2018) and IFT54 (Zhu et al., 2017) are important IFT-B interactors of kinesin 2. Interestingly, we identified two conserved amino acid patches on IFT-B that likely represent binding sites for kinesin 2 (Figure S9A). One patch is composed of IFT88/57/38 where IFT-B1 and IFT-B2 connect, while the other patch is contributed by the tetrameric CCs of IFT57/54/38/20 (Figure S9A). A *Chlamydomonas* IFT54 deletion mutant that lacks residues 342-356 no longer binds kinesin 2 *in vitro* or *in vivo* (Zhu et al., 2017). In our IFT-B model, residues 342-356 of IFT54 lie at the tetrameric interface between IFT57, -54, -38 and -20 (Figure 5C-D) and their deletion could disrupt the structure of the tetramer and thus its function as a kinesin 2 binding platform.

### Association of IFT-B with cargoes

IFT trains carry a variety of cargo into cilia including tubulin, radial spokes, and axonemal motility complexes like outer- and inner dynein arms (ODAs and IDAs) (Lechtreck et al., 2022). ODAs are imported into cilia by IFT46 via the cargo adaptor protein ODA16 (Ahmed and Mitchell, 2005; Hou et al., 2007). The N-terminal part of IFT46 (residues 1-147) interacts with ODA16 while the C-terminus is important for assembly of IFT-B complex (Hou and Witman, 2017; Taschner et al., 2017; Wang et al., 2020). The N-terminal 187 residues of IFT46 are not included in our structural models but are located at the periphery of the IFT-B complex opposite to the IFT-A binding site and are free to engage binding partners such as ODA16 (Figure 9 and S9). Residues 147-187 of IFT46 do not interact with IFT81/74, IFT52 or ODA16 and are thus free to engage other factors such as IFT56, which is implicated in the ciliary import of certain IDAs (Ishikawa et al., 2014; Xin et al., 2017).

*In vitro* studies have shown that the CH-domain of IFT81 and a highly basic amino acid stretch of IFT74N bind αβ-tubulin and likely constitutes the main αβ-tubulin cargo binding site in IFT trains (Bhogaraju et al., 2013). Although most of the αβ-tubulin that is required for axonemal growth enters cilia by diffusion (Craft Van De Weghe et al., 2020), mutations in the CH-domain of IFT81 or in IFT74N, while not hampering IFT in general, result in reduction of the frequency of anterograde IFT of αβ-tubulin to levels of 26% and 11%, respectively, when compared to control *Chlamydomonas* cells (Kubo et al., 2016). Cells with mutations in both the IFT81 CH-domain and IFT74N have almost no flagellar assembly highlighting the importance of these domains in ciliogenesis (Kubo et al., 2016). When anterograde IFT trains reach the tip of cilia, αβ-tubulin cargo must be released for the incorporation into the growing axoneme of the cilium. The release of IFT cargo such as αβ-tubulin likely employs mechanisms that weaken the interaction with IFT-trains. This hypothesis is supported by recent a study in *C. elegans,* which showed that the N-terminus of IFT74 undergoes phosphorylation by the DYF-5/MAK kinase (Jiang et al., 2022). Interestingly, the phosphorylation of IFT74N translates into a sixfold reduction in the binding affinity for αβ-tubulin (Jiang et al., 2022), which likely help facilitate the unloading of αβ-tubulin cargo from IFT trains. Interestingly, the conformation of the CH-domain of IFT81 in the second model of IFT-B presented in Figures 4E and 8B and observed in the structure of anterograde IFT trains (Lacey et al., 2022) is not compatible with the canonical association of CH domains with αβ-tubulin. It is possible that αβ-tubulin associates with both N-termini of IFT74 and IFT81 at the base of the cilium but that packaging into the anterograde IFT trains results in a re-positioning of the IFT81 CH-domain and dissociation from αβ-tubulin leaving only IFT74N attached. The final dissociation of αβ-tubulin from anterograde IFT trains may then rely on IFT74N phosphorylation as suggested by Jiang et al. 2022.

### Structural mapping of ciliopathy variants onto IFT-B

Ciliopathy variants in IFT-B proteins (Braun and Hildebrandt, 2017) were obtained from the Uniprot database and mapped onto the structure model of the IFT-B complex in Figure 9B. For this purpose, we obtained all ciliopathy related data from Uniprot, which comprises 327 distinct variants in the discussed IFT-B genes. The variants are found together with over 15 prominent ciliopathies such as Bardet-Biedl syndrome, short-rib thoracic dysplasia and asphyxiating thoracic dystrophy.

Ciliopathy variants are distributed along the IFT-B complex structure with a pronounced clustering of skeletal ciliopathy variants in IFT80 and IFT172 at the IFT-B2 end of the complex (Figure 9B). These mutations are associated with ciliopathies such as Joubert syndrome, Jeune thoracic dystrophy, asphyxiating thoracic dystrophy, short-rib thoracic dysplasia (Beales et al., 2007; Halbritter et al., 2013). The short rib polydactyly syndrome and Jeune thoracic dystrophy represent so-called ciliary chondrodysplasias, with overlapping skeletal and extra-skeletal presentations (Antony et al., 2021). Interestingly, these ciliopathies are also caused by variants that map to dynein 2 and are associated with defective retrograde IFT trains resulting in ciliary accumulation of proteins (Antony et al., 2021). This suggests that variants in IFT80 and IFT172 may cause skeletal ciliopathies by affecting the loading of IFT dynein cargo onto anterograde IFT trains.

IFT52 constitutes the backbone of IFT-B complex (Figure 8A) (Taschner et al., 2014), and IFT52 variants destabilize anterograde IFT complex assembly and disrupt ciliogenesis (Zhang et al., 2016). We have mapped 3 ciliopathy mutations associated with short rib thoracic dysplasia and one mutation associated with short rib polydactyly on or in the vicinity of the N-terminal GIFT domain of IFT52 (Figure 9B, blue and red spheres, respectively)(Chen et al., 2018; Dupont et al., 2019; Girisha et al., 2016). These ciliopathy mutations are located close to residues K130 and R204 of IFT52, where mutation to glutamate significantly reduce the IFTB1-B2 interaction (Taschner et al., 2016) and may thus destabilize IFT-B integrity (Figure 9B).

The occurrence of Bardet-Biedl syndrome (BBS) ciliopathy characterized by obesity, polydactyly, retinal degeneration, and mental retardation is typically caused by mutations or knockouts of genes that translates into proteins of the BBSome complex (Forsyth and Gunay-Aygun, 1993; Nachury et al., 2010). The BBSome complex functions as an IFT adaptor that removes membrane proteins from cilia (Lechtreck et al., 2013, 2009). Within the IFT-B complex, the IFT27/25 hetero-dimer was shown to be involved in the regulation of ciliary export of BBSomes and associated cargoes (Eguether et al., 2014; Keady et al., 2012; Liew et al., 2014). Interestingly, mutations or knockouts of IFT27/25 mimic phenotypes associated with BBS (Aldahmesh et al., 2014; Yan and Shen, 2021) or cause foetal lethality (Quélin et al., 2018). In our model, IFT27/25 is placed at the periphery of the IFT-B complex opposite to the binding site for IFT dynein cargo (Figure 9A). Given the structural flexibility between N- and C-terminal halves of IFT81/74, IFT27/25 could easily be positioned close to the ciliary membrane for BBSome interaction. Four IFT27 variants reported to be associated with BBS map to the interface with IFT74 in our model (Figure 9B, black spheres)(Aldahmesh et al., 2014; Schaefer et al., 2019). These variants could disrupt the interaction interface weakening association of IFT27 with IFT74. Recent studies support this notion, as truncations of the C-terminal region of IFT74, guided by reported missense mutations, abolish the interaction with IFT27 (Zhou et al., 2022). Improper or weakened binding of IFT27/25 on IFT81/74 is thus likely a cause for BBS.

## Materials and Methods

### Purification and reconstitution of *Chlamydomonas* IFT-B complexes

The IFT-B subcomplexes used for crosslinking/MS (Figure 1) were purified according to previously published protocols (Taschner et al., 2016, 2014; Taschner and Lorentzen, 2016). IFT80, IFT172 and the IFT46/52 complex used in interaction studies were purified according to the protocols in (Taschner et al., 2018, 2016; Wang et al., 2018).

The recombinant IFT81_460-C_/74_460-C_/27/25_1-136_ protein complex (Figure 4C-D) was obtained by co-transforming the plasmids pEC-A-His_(6)_-TEV-IFT81_460-C_, pEC-K-His_(6)_-TEV-CrIFT74_460-C_, pEC-S-His_(6)_-TEV-CrIFT25_1-136_ and pEC-Cm-CrIFT27 into *E. coli* BL21 (DE3) cells. The IFT81_534-623_/74_533-615_/27/25_1-136_ complex (Figure 4C-D) was also produced by co-transformation of the respective plasmids in *E. coli* (DE3). IFT81_460-C_/74_460-C_/27/25_1-136_ or IFT81_534-623_/74_533-615_/27/25_1-136_ were over-expressed in cultures of 6L of Terrific broth (TB) medium supplemented with the appropriate antibiotics. The bacterial cultures were grown at 37°C until OD_600_ reached 0.5, cooled down to 18°C and induced with 0.5mM of Isopropyl β-D-1-thiogalactopyranoside (IPTG) for 18 hours to trigger the expression of recombinant proteins. The cultures were harvested by centrifugation (rotor F9-6x1000lex, at 7822 RFC (Relative Centrifugal Force), 4°C for 12min), typically yielding 200g of wet cell pellet. The cell pellets were dissolved in 200ml of lysis buffer (50mM Tris pH 7.5, 150mM NaCl, 10% (v/v) glycerol, 1mM MgCl_2_ and 5mM βME) supplemented with 2 tablets of cOmplete EDTA free protease inhibitor, 0.5mM of phenylmethylsulfonyl fluoride (PMSF) and DNase 1 (1U/µl) prior to cell lysis by sonication. The cell lysate was cleared by centrifugation at 30000 x g for 30 min and the supernatant was collected, filtered through 5µm filters and circulated through a pre-equilibrated 5ml cOmplete Ni^2+^-NTA column using a peristaltic pump. The column was further washed with lysis buffer containing 20mM Imidazole pH 7.5, high salt buffer (50mM Tris pH 7.5, 1M NaCl, 10% (v/v) glycerol, 1mM MgCl_2_ and 5mM βME) and low salt buffer (50mM Tris pH 7.5, 75mM NaCl, 10% (v/v) glycerol, 1mM MgCl_2_ and 5mM βME). A 5ml HiTrap Hp Q anion column equilibrated with low salt buffer was mounted below the Ni^2+^-NTA column and the protein complexes were eluted from both columns in 5 elution steps each with 25ml elution buffer (50mM Tris pH 7.5, 75mM NaCl, 10% (v/v) glycerol, 600mM imidazole, 1mM MgCl_2_ and 5mM βME). The elutions were concentrated to 1ml and loaded onto a HiLoad 16/600 Superdex 200 (GE Healthcare) column equilibrated in SEC buffer (10mM HEPES pH 7.5, 150mM NaCl, 1mM MgCl_2_ and 1mM DTT). The fractions containing pure protein complexes were pooled, concentrated, snap cooled in liquid nitrogen and stored at −70°C until use.

### Protein complex prediction with AlphaFold multimer

For predicting the structure of IFT-B sub-complexes we used a modified version of AlphaFold v2.1.0 on Colab notebook for protein complexes smaller than 1200 residues (Mirdita et al., 2022) as well as a local installation of AlphaFold multimer for larger sub-complexes (Evans et al., 2022; Jumper et al., 2021). AlphaPickle was used to extract the predicted alignment score from AlphaFold runs (mattarnoldbio, 2021). The structural model of the 15 subunit IFT-B complex was assembled from the structural models of smaller modules using the relevant *Chlamydomonas reinhardtii* proteins sequences. All sequences used for structure prediction have at least 500 homologs in available sequence databases and all structural predictions shown in the figures have low PAE scores for the interacting regions indicating a high degree of certainty in the relative positions of subunits within the complexes. A total of 9 sub-complexes were predicted using AF and subsequently assembled into the 15 subunit IFT-B complex in PyMOL v. 2.5 (Schrodinger LLC, https://pymol.org) using the align function. The long regions with pLDDT scores lower than 50 are predicted to be unstructured and were excluded from the assembled model of 15 subunit IFT-B complex shown in Figures 8–9.

### Site directed photo-crosslinking

To incorporate pBpa into *Chlamydomonas* IFT81 we replaced the native DNA sequence at the desired position (E418 or F68) with the TAG sequence that encodes pBpa. As IFT81 can be produced only as part of the IFT81/74/27/25 complex, we co-transformed the pEC-A-His_(6)_-TEV-IFT81, pEC-K-His_(6)_-TEV-CrIFT74, pEC-S-His_(6)_-TEV-CrIFT27-RBS-CrIFT25_1-136_ plasmids carrying the IFT genes together with the pEVol suppression plasmid (Young et al., 2010) into *E. coli* BL21 (DE3) cells. The bacterial cultures were grown in TB media supplement with 0.5mM pBpa, at 37°C until OD_600_ reached 0.5, cooled down to 18°C and induced with 0.5mM IPTG and 0.2% L-arabinose for 18 hours to trigger the expression of recombinant IFT proteins as well as the production of pBpa-tRNA. In total, 6L of pBpa cultures were grown simultaneously with 1L of IFT22 culture and harvested together to yield a pentameric IFT81/74/27/25/22 complex. Subsequently the protein complex was purified as previously published (Taschner et al., 2011) except that the UV lamp was turned off during SEC to prevent the premature activation of pBpa. Samples containing 7µM of purified IFT81/74/27/25/22 complexes with or without pBpa were exposed 20 minutes to 365nm wavelength UV light (UV chamber: BLX-365 from Vilber Lourmat) for crosslinking. 8µl from each sample were denatured, loaded onto SDS-PAGE and the resulting gel was stained with Coomassie brilliant blue.

### Rigid-body assembly of the 15-subunit *Chlamydomonas* IFT-B complex

To assemble the IFT-B complex, the predicted structure of IFT81/74_120-C_/27/25_1-136_/22 (Figure 2) was aligned onto the C-termini of IFT81 and IFT74 of the predicted structure of IFT88_120-713_/70/52/46_188-319_/81_587-645_/74_583-C_ with an RMSD of 1Å. Next, the IFT88_120-713_/52_1-336_/57_356-C_/38_174-303_ structural model was docked onto the IFT88_120-713_/70/52/46_188-319_/81/74_120-C_/27/25_1-136_/22 complex by superimposition of the IFT88 subunit yielding an RMSD of 3Å. The N-terminal globular domain of IFT52 aligned well (although it was not explicitly used in the superpositioning) and preserved its position on the central part of C-termini of IFT57 and IFT38. In the following step, the structure of IFT57/38/54_135-C_/20 was added to the model by superpositioning onto IFT57_358-469_/38_173-303_ (RMSD of 2Å). The predicted IFT80/38 structure had the IFT38 C-terminal domain removed because of low pLDDT scores in absence of its interacting partner IFT57. The remaining model, including full length IFT80 and the CH-domain of IFT38, was docked onto the IFT88_120-713_/70/52/46_188-319_/81/74_120-C_/27/25_1-136_/22/57/38/54/20 complex via the CH-domain of IFT38 (RMSD of 0.2Å), which resulted in a structural model of the 14 subunit IFT88_120-713_/70/52/46_188-319_/81/74_120-C_/27/25_1-136_/22/57/38/54/20/80 complex. Finally, the structural model of IFT172_1-785_/80 (Figure S6E panel A) was docked onto IFT80 of the IFT-B 14mer (RMSD of 1Å) and the CH-domain ofIFT57 added via superpositioning of the IFT172/57CH model (RMSD of 0.8Å) yielding a structural model of the 15 subunit IFT-B complex as shown in Figure 8. The final model was subsequently subjected to 5 macrocycles of geometric energy minimization in PHENIX (Adams et al., 2010) using geometric, non-bonding and secondary structure restraints.

### Small angle X-ray scattering (SAXS) measurements of CrIFT80/38 complexes

The SAXS experiment on IFT80/38 shown in Figure 6D were performed at the BM29 beamline (ESRF, Grenoble, France) using a Pilatus 1M detector using the SEC/SAXS protocol described in (Brennich et al., 2017). SAXS data were collected on the purified IFT38_1-133_ protein or the IFT80_1-582_/38_1-133_ complex eluting directly from a Superdex 200 10/300 GL column. Correction for radiation damage, data merging and buffer subtraction were performed on site. We used GNOME to extract SAXS parameters such as maximum particle size (Dmax) (Svergun, 1992) from the ATSAS package software (Petoukhov et al., 2012). The theoretical SAXS curve of IFT80_1-582_/38_1-133_ was calculated from the structural model using CRYSOL (Svergun et al., 1995) and fitted to the experimental data.

### Mass spectrometry analysis of DSBU crosslinked protein complexes

Cross-linked proteins were digested to generate peptides for MS-analysis. Initially, proteins were reduced and alkylated in a buffer containing 5% SDS, 10mM TCEP and 11mM CAA in 100mM Tris-HCl pH 8.5 for 10 minutes at 95^0^C and precipitated on MagResyn HILIC magnetic particles (Resyn Biosciences) in 70% acetonitrile for 20 minutes and washed with 95% acetonitrile and 70% ethanol before on-bead digestion with Lys-C and trypsin in 50mM ammonium bicarbonate pH 8.0 overnight (Batth et al., 2019). The resulting peptides were desalted on Sep-Pak tC18 cartridges (Waters Corporation) and subjected to direct MS-analysis (10% of the sample) or further enriched for multiple charged cross-linked peptides on mixed-mode C18/SCX cartridges (Oasis MCX, Waters Corporation) generating four fractions for MS-analysis as previously described (Iacobucci et al., 2018). Cross-linked peptides were analyzed by an Easy nanoLC system coupled to an Orbitrap Exploris 480 mass spectrometer (Thermo Scientific) using data-dependent acquisition of ions with a charge state between 3 and 8 and HCD fragmentation using stepped normalized collision energy (NCE) of 27, 30 and 33. Interpeptide cross-links were identified based on their signature doublet signals from the cleavable DSBU cross-linker as implemented in the RISEUP mode of the Program MeroX version 2.0 (Götze et al., 2019). The identified cross-links were evaluated by scores for peptide pair identifications and cross-link position assignments within the peptide sequences.

### Labeling of crosslinking pairs onto the structural models

The MS/crosslinking results containing residue to residue intra- and intermolecular crosslinks with scores >80 of a recombinant *Chlamydomonas* IFT-B1 nonameric complex (IFT88_1-437_/70/52_281-430_/46_188-319_/81/74_128-C_/27/25_1-136_/22) and of an IFT-B1_B2 hexameric complex (CrIFT88/52_1-335-GST_/57/38/54/20 complex) were used. AlphaFold predicted sub-structures were used as scaffold, but regions predicted to be disordered (containing pLDDT scores <50) were discarded from the analysis.

Python 3.8 was used to generate a list of commands drawing all the crosslinking labels simultaneously. As inputs were used the protein-to-chain conversion map, the IFT-B 15mer in pdb format containing the positions of atoms and the amino acid pairs resulting from mass-spectrometry. The distances between residue pairs within the IFT-B 15mer were calculated and classified according to their length as short (< 32Å) or long (>32Å) range crosslinking pairs. The threshold was chosen based on a previous report, which identified DSBU crosslinking pairs at a reliable distance of 27Å (Felker et al., 2021). We decided to increase the threshold to 32Å to account for a predicted alignment error (PAE) of 5Å in the AF models. For validation of the automatic labelling procedure, comparison was done with manually labelling using the Wizard tool of PyMOL v. 2.5.

### Pull-down experiments

For CrIFT57-CrIFT172 pulldown assays, GST-tagged CrIFT57_1-234_ or free GST tag was immobilized onto GSH resin by incubating 10µM GST-IFT57_1-234_ or GST-tag in 100µL buffer B1 (10mM HEPES pH 7.5, 100mM NaCl, 5% Glycerol and 1mM DTT) with 30 µL bed volume of GSH resin for 1h at 4°C. After incubation, the beads were washed 3 times with Buffer B1. CrIFT172_1-968_ was diluted to 20µM in Buffer B1 and 100µL of this sample was incubated with the prepared GSH resin (loaded with GST-tag or GST-IFT57_1-234_) for 2h at 4°C. The beads were washed 3 times with Buffer B1, and bound proteins eluted by incubating the beads with Buffer B1 containing 30mM reduced glutathione. 3μg of proteins were loaded as input.

For the CrIFT80-CrIFT172 pulldown assays, Venus-tagged CrIFT80 was immobilized onto GFP-binder bead resin by incubating 5µM CrIFT80-Venus in 100µL buffer B1 with 20µL bed volume of GFP binder bead resin for 1h at 4°C. A control of unbound beads was included. After incubation, the beads were washed 2 times with buffer B1. CrIFT172 or CrIFT172_968-C_ was diluted to 15µM in buffer B1 and 100µL of this sample was incubated with the CrIFT80-Venus preloaded GFP-binder bead resin (or unbound beads for control) for 2h at 4°C. After incubation, beads were washed 4 times with buffer B1, and bound proteins were eluted by incubating the beads with 0.1M citric acid. 3μg of proteins were loaded as input.

### X-ray diffraction analysis of CrIFT88_1-437_/70/52_281-360_ and CrIFT70/52_330-430_/46_165-319_ complexes

CrIFT88_1-437_/70/52_281-360_ was crystallized by mixing equal volumes of the complex (buffer: 20 mM Tris pH 8.0, 450mM NaCl, 7.5% glycerol and 2.5mM DTT) at a concentration of 33mg/ml with a crystallization solution of 0.1M Tris pH 8.5 and 25% PEG 6k using the sitting drop vapor diffusion method. The CrIFT70/52_330-430_/46_165-319_ complex (buffer: 10mM Hepes pH 7.5, 150mM NaCl and 2 mM DTT) at 25 mg/ml was mixed with an equal volume of 50mM Tris pH 8.0 and 8% PEG3350 for crystallization. Native X-ray diffraction data were collected at the Swiss Light Source (SLS; Villigen, Switzerland) at the PXII beamline on a Pilatus 6M detector. Crystals of either complex diffracted to about 4Å resolution and complete datasets were collected and processed with XDS (Kabsch, 2010) and AIMLESS as part of the CCP4 package (Winn et al., 2011). Molecular replacement was carried out using the CrIFT70/52 crystal structure previously published (Taschner et al., 2014) and AF models of CrIFT52C/46C and IFT88 in the program Phaser (Storoni et al., 2004). Data from both crystal forms were originally processed in orthorhombic space groups but the true crystal systems were determined as monoclinic during molecular replacement and refinement. Following molecular replacement, the models were manually rebuilt in Coot (Emsley et al., 2010) removing structural elements without clear electron density followed by refinement in PHENIX (Adams et al., 2010). Data and refinement statistics are listed in Supplementary Table 1.

### Molecular dynamic flexible fitting (MDFF)

The model corresponding to the IFT-B 15mer was manually fitted into the cryo-ET density of IFT-B trains as rigid bodies using the phenix.dock_in_map feature of the PHENIX package (Adams et al., 2010). The approximate rigid-body fit was used as input for the Namdinator (Kidmose et al., 2019), a locally installed molecular dynamic tool for flexible fitting. After 400.000 reiterations, the resulting models were re-fitted in the cryo-ET density and Namdinator was run for another 400.000 cycles to reach final fit. The resulting IFT-B models were subsequently real-space refined in PHENIX (Adams et al., 2010) to produce the final model.

### Mapping of single point mutation causing ciliopathies onto the IFT-B 15mer structure

The single-residue variant information for IFT-B proteins was extracted from the Uniprot variants file homo_sapiens_variation.txt.gz (retrieved on 02.08.2022). Information was pre-processed before aligning the human proteins according to their accession number onto the *Chlamydomonas reinhardtii* sequences used for modeling in this study. Eight of the 436 variants did not have a corresponding *Chlamydomonas reinhardtii* residue in the alignment and were thus mapped to the closest preceding amino acid. Disease groupings were introduced according to the first two words in the disease name, specifically to group Asphyxiating thoracic dystrophy, Bardet-Biedl Syndrome, Retinitis Pigmentosa and Short-rib thoracic dysplasia. Number of variants associated with every group was counted and those groups/phenotypes associated with more than 3 variants were plotted with separate colors in the model. All the remaining variants are shown plotted in red. The spheres display the C-alpha atoms of the variants.

## Supporting information

Supplementary data for crosslinking of IFT-B1

Supplementary data for crosslinking of IFT-B1-B2

Supplemental information for ciliopathy variants used in Figure 9B

Movie showing structural mapping of IFT25-IFT81/74 crosslinks

Movie showing structural mapping of IFT27-IFT81/74 crosslinks

Movie showing structural mapping of IFT88/70-IFT52/46 crosslinks

Movie showing structural mapping of IFT38-IFT57-52 crosslinks

Movie showing structural mapping of IFT88-IFT38-IFT52 crosslinks

## Acknowledgements

We thank Kathrine Kjærgaard Frederiksen and Anni Christensen for technical assistance with cloning and/or protein purification and Jesper Lykkegaard Karlsen for computational assistance with running Alphafold and Namdinator. We also thank Hugo van den Hoek and Benjamin D. Engel for making their unpublished cryo-ET maps available to us. This work was funded by grants from the Novo Nordisk Foundation (grant number NNF15OC00114164) and the Independent Research Fund Denmark (grant no: 1026-00016B) to E.L. N.A.P was supported by a postdoc fellowship from the European Commission (H2020, Grant Agreement number 888322). R.B.R. and E.L. have received funding from the European Union’s Horizon 2020 research and innovation programme under the Marie Skłodowska-Curie grant agreement No. 861329. M.L.L and J.S.A was supported by the Independent Research Fund Denmark (grant no: 8021-00425B) to J.S.A.

