## Supplemental figures and tables for "Biochemically validated structural model of the 15-subunit IFT-B complex"

Supplementary Data
for

### Biochemically validated structural model of the 15-subunit IFT-B complex

Narcis A. Petriman[1*], Marta Loureiro-López[2], Michael Taschner[3], Nevin K. Zacharia[1], Magdalena M. Georgieva[4], Niels Boegholm[1], André Mourão[5], Robert B. Russell[4], Jens S. Andersen[2] and Esben Lorentzen[1*]

[1] Department of Molecular Biology and Genetics, Aarhus University, Universitetsbyen 81, 8000 Aarhus C, Denmark

[2] Department for Biochemistry and Molecular Biology, University of Southern Denmark, Campusvej 55, 5230 Odense M, Denmark

[3] Department of Fundamental Microbiology, University of Lausanne, CH-1015 Lausanne, Switzerland

[4] BioQuant, Heidelberg University, Im Neuenheimer Feld 261, 69120, Heidelberg, Germany

[5] Institute of Structural Biology, Helmholtz Zentrum München, Ingolstädter Landstrasse 1, 85764 Neuherberg, Germany

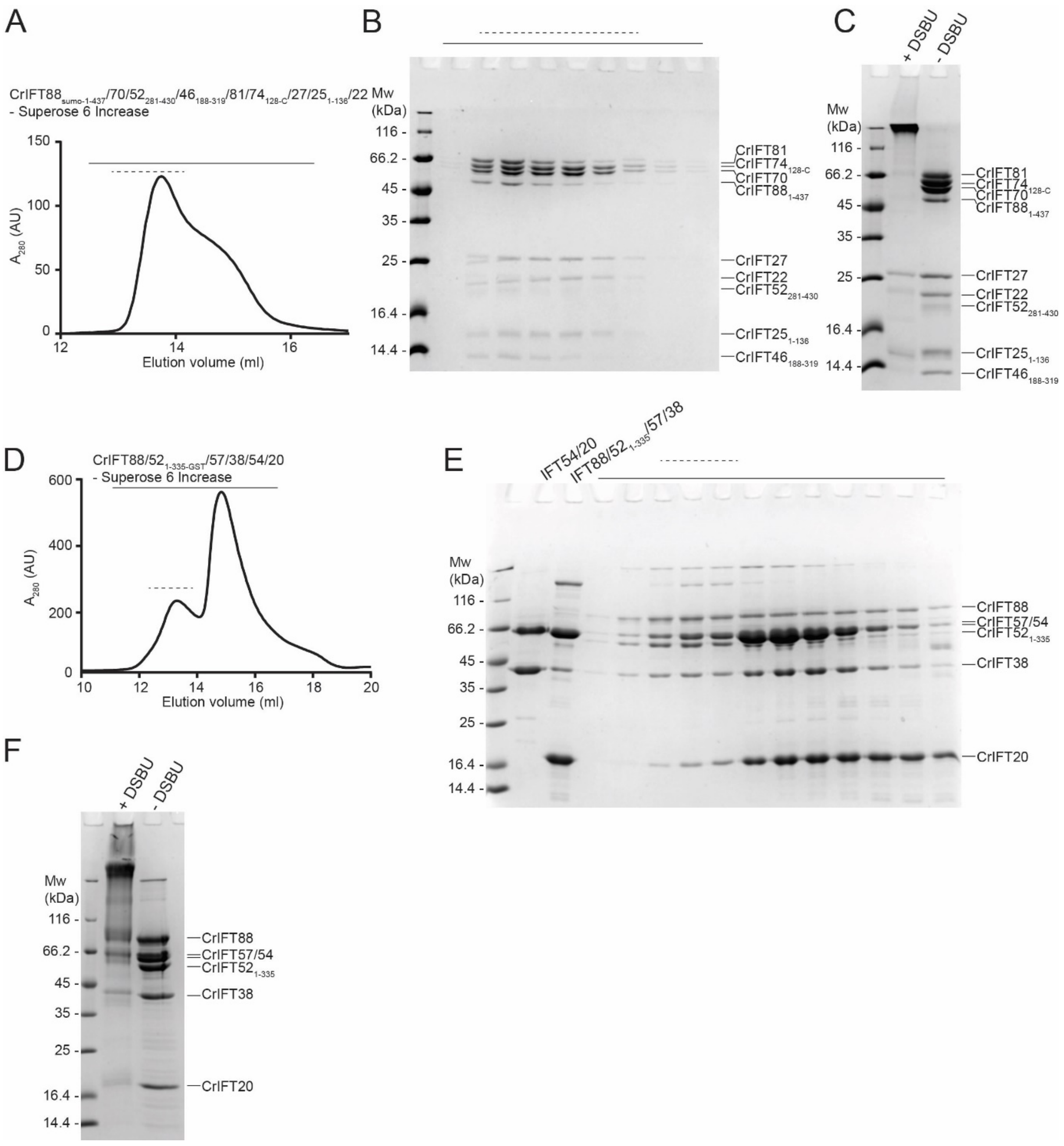


**Figure S1: Purification and crosslinking of CrIFT88$_{sumo\text{-}1\text{-}437}$/70/52$_{281\text{-}430}$/46$_{188\text{-}319}$/81/74$_{128\text{-}C}$/27/25$_{1\text{-}136}$/22 and CrIFT88/52$_{1\text{-}335\text{-}GST}$/57/38/54/20 complexes**

**(A)** SEC profile of purified CrIFT88$_{sumo\text{-}1\text{-}437}$/70/52$_{281\text{-}430}$/46$_{188\text{-}319}$/81/74$_{128\text{-}C}$/27/25$_{1\text{-}136}$/22 complex. **(B)** All fractions illustrated above the SEC profile in panel (A) with a solid horizontal line are verified for purity on SDS PAGE and stained with Coomassie. The dashed lines in (A) and (B) represent the SEC fractions that were pooled for crosslinking. **(C)** The pooled SEC fractions were crosslinked with 0.25mM DSBU and subjected to MS analysis. **(D)** SEC with incubated CrIFT54/20 and CrIFT88/52$_{1\text{-}335\text{-}GST}$/57/38 complexes to yield a CrIFT88/52$_{1\text{-}335\text{-}GST}$/57/38/54/20 hexameric complex. The SEC fractions highlighted with a solid line were monitored on SDS PAGE for purity (E) and fractions containing all 6 subunits indicated by a dashed line were pooled, crosslinked as shown in (F) and subjected to MS analysis.

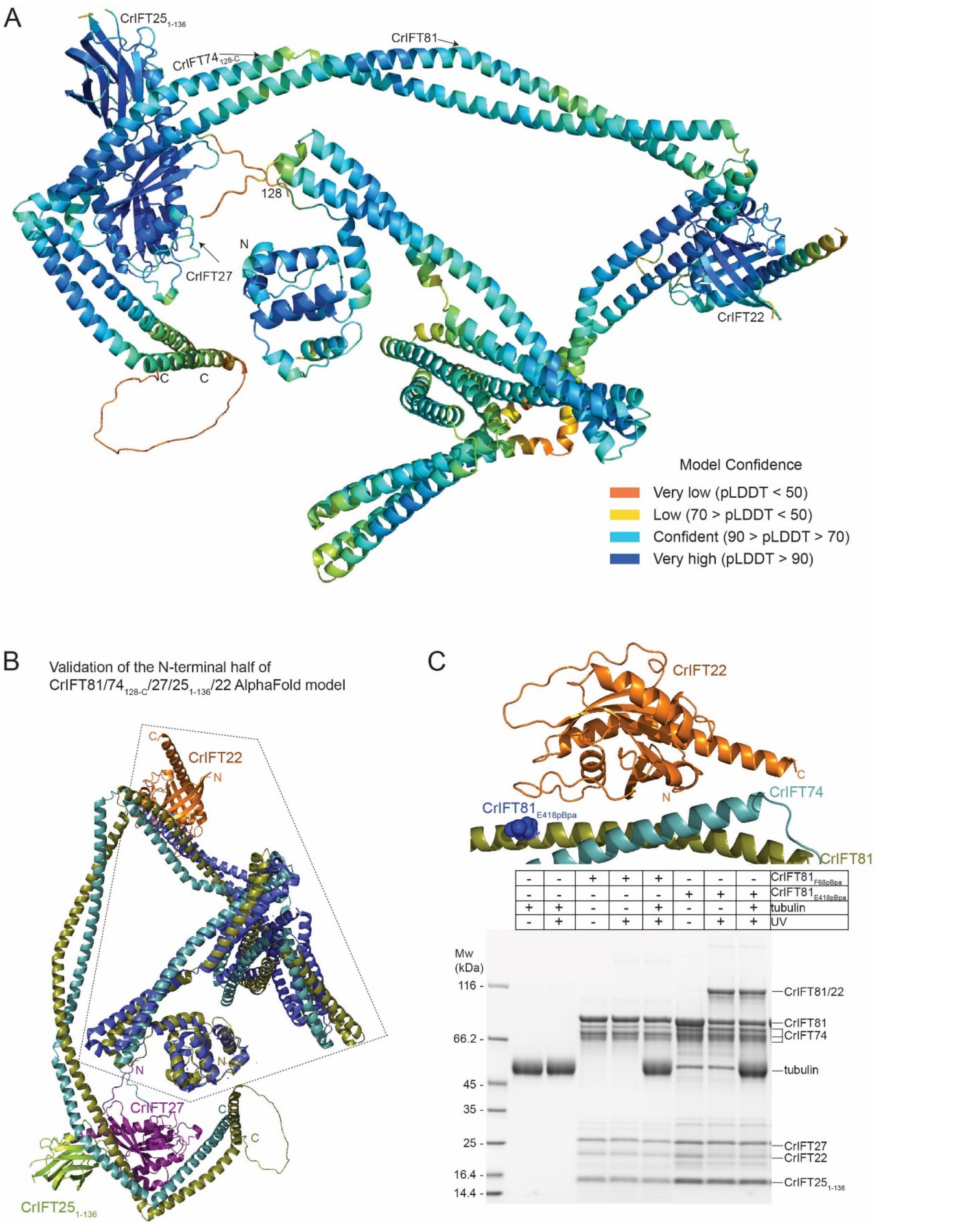


**Figure S2: AF model of IFT81/74/27/25/22 and validation of the IFT22 binding site**

**(A)** AlphaFold predicted structure of CrIFT81/74$_{128\text{-}C}$/27/25$_{1\text{-}136}$/22 colored according to pLDDT scores.

**(B)** The N-terminal half of the predicted CrIFT81/74$_{128\text{-}C}$/27/25$_{1\text{-}136}$/22 model superimposed onto the crystal structure of TbIFT81N/74N/22 (PDB accession number 6ian; colored blue). **(C)** The position of CrIFT22 in the predicted AlphaFold model is validated by site directed photo-crosslinking. The recombinant CrIFT81/74/27/$25_{1-136}$/22 complex containing acid p-benzoyl-L-phenyl-alanine (pBpa) instead of its natural E418 amino acid in CrIFT81 was activated by UV light (365nm wavelength) and showed a strong CrIFT81-CrIFT22 crosslink as illustrated on SDS PAGE and Coomassie staining (last two lanes). The CrIFT81-CrIFT22 crosslink formed at the interaction interface predicted by AlphaFold and was independent of tubulin cargo, which binds to the N-termini of CrIFT81/74. As a negative control, we used a recombinant CrIFT81/74/27/$25_{1-136}$/22 complex that had the F68 residue located at the N-terminus of CrIFT81, far from the CrIFT22 binding site, replaced by pBpa (lanes 3 - 5).

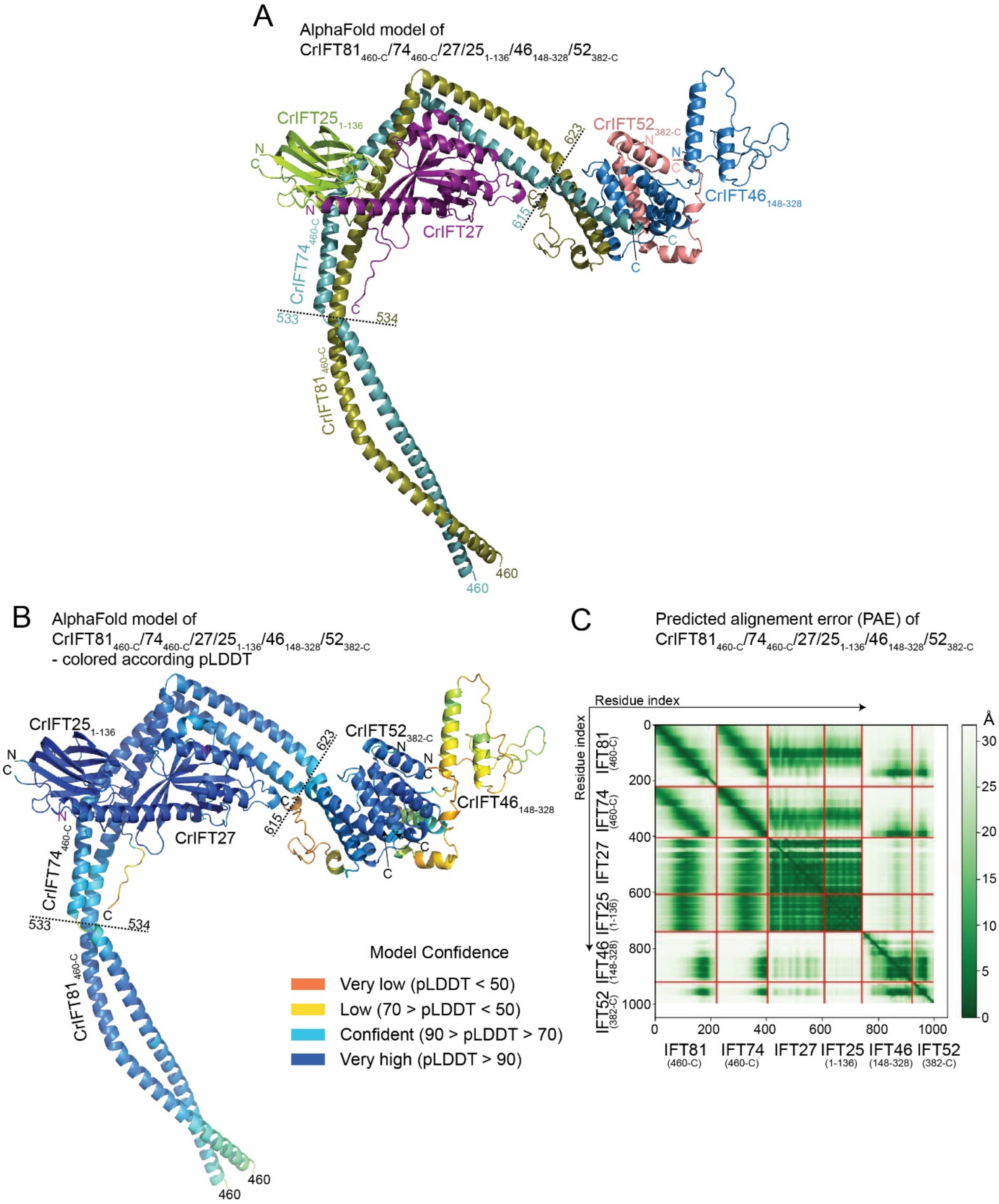


**Figure S3: AF model of IFT81/74/27/25/46/52 complex**

(**A**) The AF structure prediction of CrIFT81$_{460\text{-}C}$/74$_{460\text{-}C}$/27/25$_{1\text{-}136}$/46$_{148\text{-}328}$/52$_{382\text{-}C}$ complex colored according to chains. (**B**) The CrIFT81$_{460\text{-}C}$/74$_{460\text{-}C}$/27/25$_{1\text{-}136}$/46$_{148\text{-}328}$/52$_{382\text{-}C}$ structure is predicted with very high confidence (pLDDT scores >70 except for less ordered termini) and illustrates the docking of CrIFT27/25$_{1\text{-}136}$ heterodimer and CrIFT52$_{382\text{-}C}$/46$_{148\text{-}328}$ on the C-terminal half of CrIFT81/74. **(C)** The PAE plot of CrIFT81$_{460\text{-}C}$/74$_{460\text{-}C}$/27/25$_{1\text{-}136}$/46$_{148\text{-}328}$/52$_{382\text{-}C}$ complex.

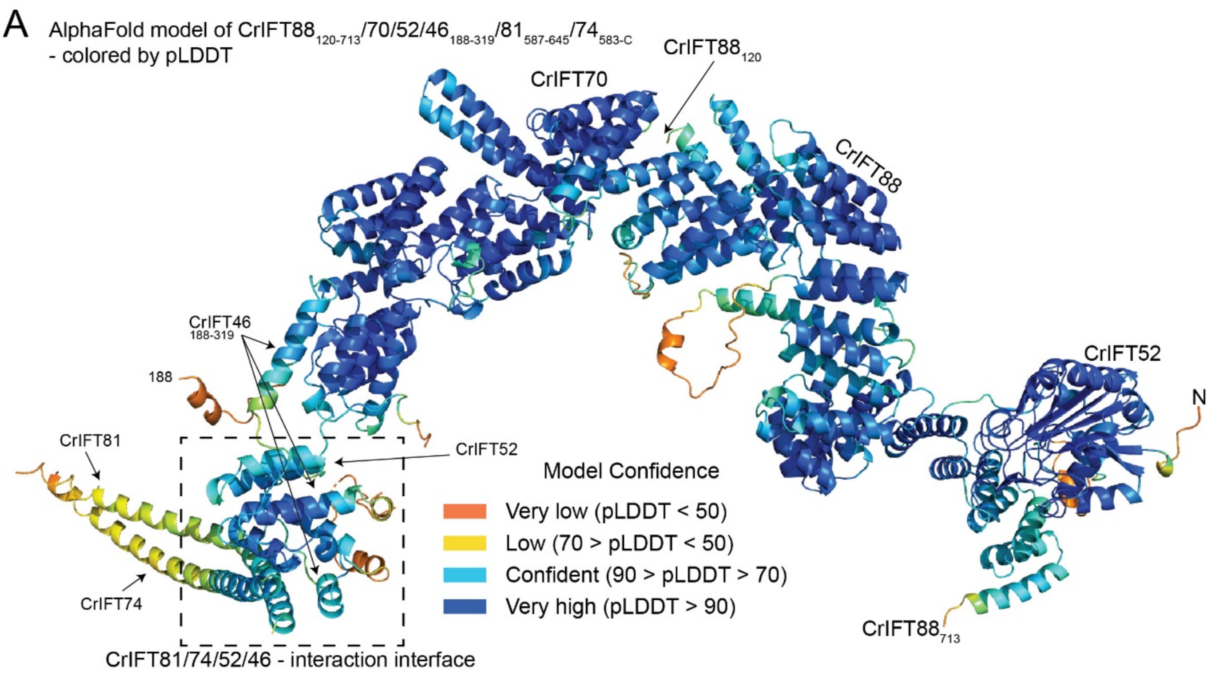


**Figure S4: (A) Alphafold predicted model of IFT88/70/52/46 with IFT81/74 C-terminal helices**

The predicted structure of the CrIFT88$_{120\text{-}713}$/70/52/46$_{188\text{-}319}$/81$_{587\text{-}645}$/74$_{583\text{-}C}$ complex colored according to the pLDDT confidence score. The lower pLDDT score for part of the IFT81/74 helices is likely a result of the short constructs of IFT81/74 and the missing IFT27/25 subunits in this prediction.

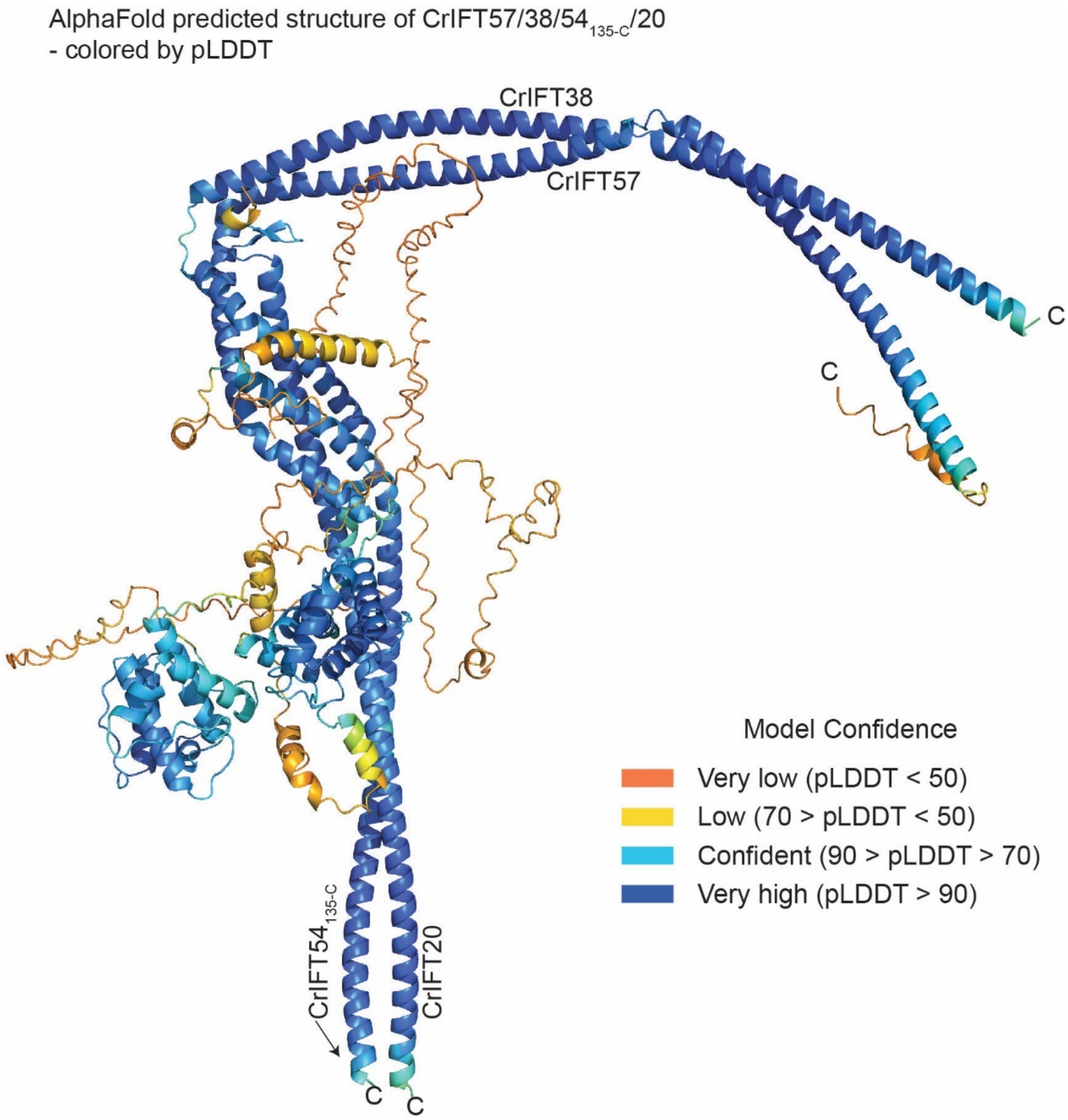


**Figure S5: AlphaFold predicted model of the CrIFT57/38/54$_{135\text{-}C}$/20 tetramer colored according the pLDDT scores**

The N-terminal CH-domain of CrIFT57 as well as the coiled-coils of all four subunits are predicted with high confidence (pLDDT>70). Intrinsically disorder structural elements between CH and CC domains are colored orange and have pLDDT scores <50.

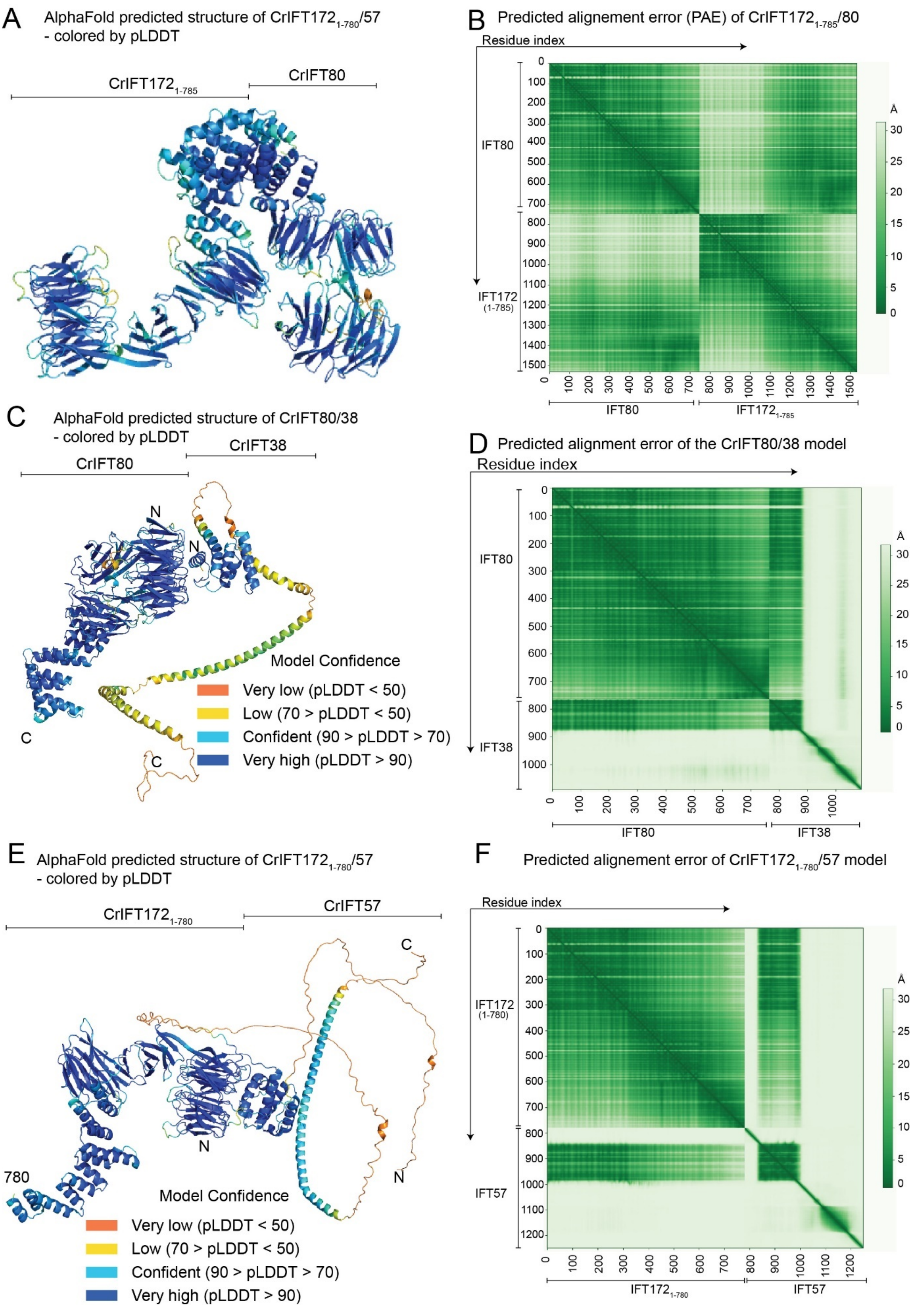


**Figure S6: AlphaFold predictions of CrIFT172/80, CrIFT172/57 and CrIFT80/38 complexes**

**(A)** The predicted structure of $IFT172_{1\text{-}785}$ in complex with IFT80 colored according to pLDDT scores. **(B)** The predicted alignment error plot for the $IFT172_{1\text{-}785}$/80 structure. **(C)** The predicted AlphaFold model of CrIFT80 and CrIFT38 colored according to the pLDDT score. Structures of CrIFT80 and the CH-domain of CrIFT38 are predicted with high-confidence scores, whereas very low confidence scores accompany the structure the C-terminal CC of CrIFT38 in the absence of its interacting partners. **(D)** The predicted alignment error plot for CrIFT80/30 model. The 1-300 indexed residues corresponding to the first β-propeller of CrIFT80 and the 748-875 indexed residues corresponding to the CH-domain of CrIFT38 have low PAE scores demonstrating high confidence for the multimer structure prediction. **(E)** The predicted structural model of $CrIFT172_{1\text{-}780}$ in complex with CrIFT57 colored according to the pLDDT scores. Very low confidence scores are observed for the linker region between the CH-domain of CrIFT57 and the helical C-terminal domain. Otherwise, the folding of the complex is predicted with high confidence. **(F)** The predicted alignment error of the $CrIFT172_{1\text{-}780}$/57. Low PAE scores are calculated between residues corresponding to $CrIFT172_{1\text{-}780}$ (indexed as 1-300) and CrIFT57 (indexed as 850-900).

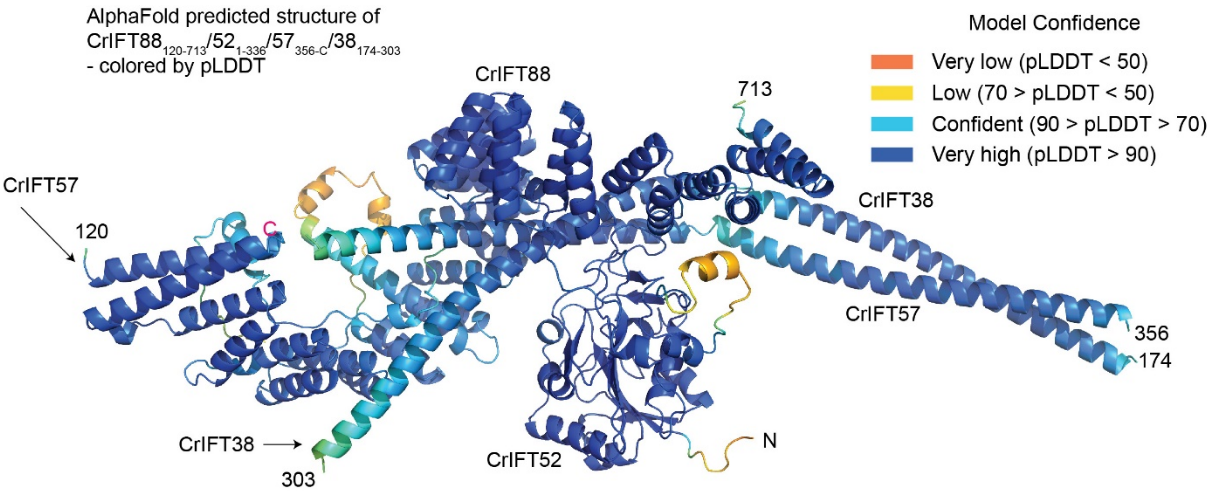


**Figure S7: IFT88 bridges the interaction between IFT-B1 and IFT-B2 complexes**

The AlphaFold predicted structure of the $CrIFT88_{120\text{-}713}/52_{1\text{-}336}/57_{356\text{-}C}/38_{174\text{-}303}$ complex colored according to the pLDDT confidence score.

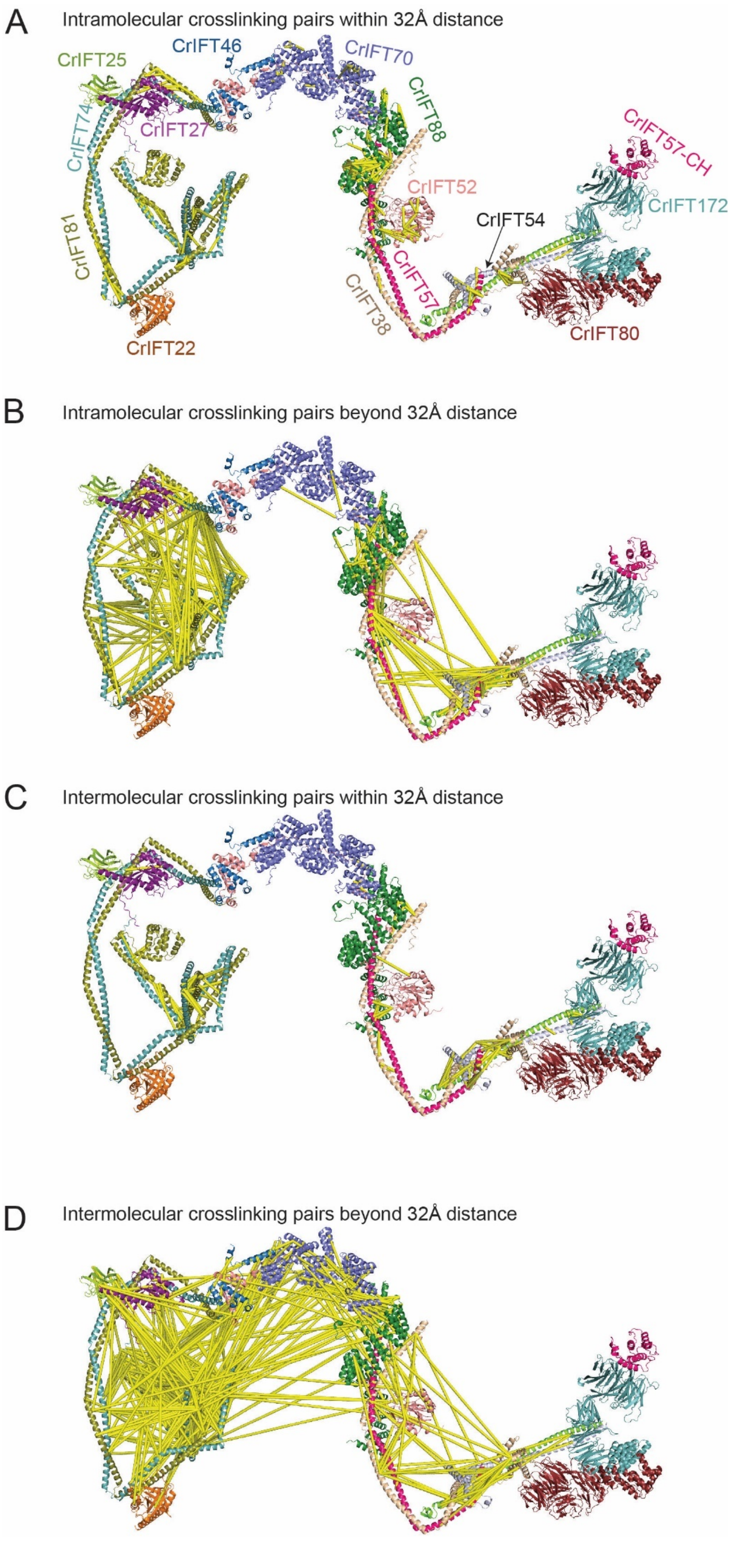
A
Intramolecular crosslinking pairs within 32Å distance
CrIFT25
CrIFT46
CrIFT70
CrIFT88
CrIFT74
CrIFT27
CrIFT81
CrIFT22
CrIFT52
CrIFT54
CrIFT57-CH
CrIFT172
CrIFT57
CrIFT38
CrIFT80
B
Intramolecular crosslinking pairs beyond 32Å distance
C
Intermolecular crosslinking pairs within 32Å distance
D
Intermolecular crosslinking pairs beyond 32Å distance

**Figure S8: Global mapping of DSBU crosslinking onto the structural model of the 15 subunits IFT-B complex**

**(A)** Complete short-range (<32Å distance) intramolecular crosslinks identified by MS mapped onto the IFT-B 15mer structure. **(B)** Long-range (>32Å distance) intramolecular crosslinks identified by MS mapped onto the IFT-B 15mer structure. **(C)** Complete intermolecular crosslinks identified by MS that fall within 32Å. **(D)** Complete intermolecular crosslinks identified by MS that fall beyond 32Å.

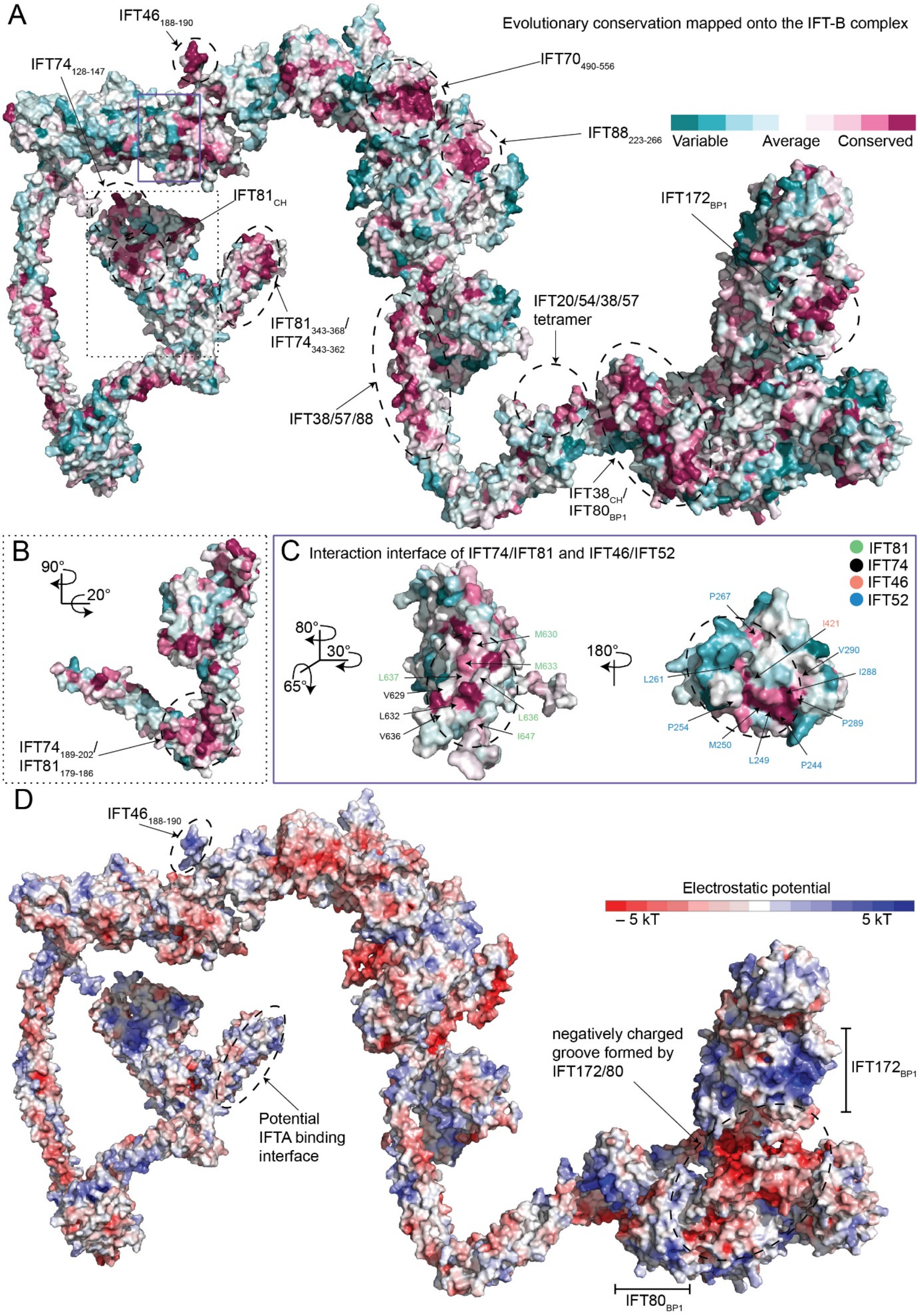


**Figure S9: Conservation and electrostatic plots of the 15-subunit IFT-B complex structure**

**(A)** Conservation analysis of the IFT-B highlights evolutionary conserved patches on the IFT-B complex. The analysis was performed with the ConSurf server (Ashkenazy et al., 2016, 2010; Celniker et al., 2013). Dashed and solid rectangles outline the regions which are discussed further in (B) and (C). **(B)** Side view of the N-terminal regions of IFT81/74 emphasizing the CH-domain of IFT81 and the conserved patch on CCs I and II where the predicted interaction with IFT88 takes place. **(C)** The conservation surface plot illustrates the interaction interface between the CC X of IFT81/74 and the C-terminal domains of IFT52/46. Conserved hydrophobic residues at the interaction interface are labeled for IFT81 (green) and IFT74 (black) in the left figure and for IFT52 (salmon) and IFT46 (blue) in the right figure. **(D)** Electrostatic plot highlighting the charged regions on IFT81/74 CC V, the platform on IFT80/172 where cargo-dynein is proposed to bind and the positively charged region on IFT46.

**Movie 1: 3D representation of the crosslinking network of CrIFT25$_{1\text{-}136}$ with IFT74$_{460\text{-}C}$ and IFT81$_{460\text{-}C}$**

The 3D representation of CrIFT25$_{1\text{-}136}$ crosslinking network is displayed on the AlphaFold predicted structure of CrIFT81/74$_{128\text{-}C}$/27/25$_{1\text{-}136}$/22 (see Fig.2A and S5). CrIFT81 is colored in olive-green, CrIFT74 in teal, CrIFT27 in magenta and CrIFT25$_{1\text{-}136}$ in lime-green. Identified crosslinks are shown between residues of IFT25 and interacting partners as dashed blue lines.

**Movie 2: 3D representation of the crosslinking network of CrIFT27 with CrIFT74$_{460\text{-}C}$ and CrIFT81$_{460\text{-}C}$**

The same color code and labelling strategy as in movie 1 is used. The 3D crosslinking network of CrIFT27 with both CrIFT81 and CrIFT74 is shown as dashed lines that connects side chains of reactive groups.

**Movie 3: 3D representation of the crosslinking network within the CrIFT70/88$_{1\text{-}437}$/52$_{281\text{-}430}$/46$_{188\text{-}319}$**

CrIFT88 is colored in green, CrIFT70 in purple, CrIFT52 in salmon and CrIFT46 in skyblue. All the crosslinking pairs are labelled as dashed blue lines connecting the sidechains of reactive residues.

**Movie 4: 3D representation of the crosslinking network of CrIFT38 with CrIFT57 and CrIFT52 with both CrIFT57 and CrIFT38 within the CrIFT88$_{120-713}$/52$_{1-336}$/57$_{356-C}$/38$_{174-303}$ complex**

CrIFT88 is colored in green, CrIFT52 in salmon, CrIFT57 in pink and CrIFT38 in beige. All the crosslinking pairs are labelled as dashed blue lines connecting the sidechains of reactive residues.

**Movie 5: 3D representation of the crosslinking network of CrIFT88 with CrIFT38 and CrIFT52 within the CrIFT88$_{120-713}$/52$_{1-336}$/57$_{356-C}$/38$_{174-303}$ complex**

The same color code and labeling strategy as in Movie 4 is used.

**Supplementary material 1: Crosslinks identified within the IFT-B1 nonamer by XL-MS/MS**

An excel file containing MeroX output after the analysis of crosslinking pairs found within the *Chlamydomonas* IFT-B1 nonamer at a high confidence based on a false discovery rate (FDR) of 1%. The "score" reflects the best score calculated for any combination of the cross-link sites in the two investigated peptides. The "site1" and "site2" represents crosslinking pairs between the first (Prot1) and the second (Prot2) protein. "P%" shows the probability of the site to be identified correctly and "type" indicates whether the crosslinks are taking place within the same protein (intraprotein) or between two different proteins (interprotein).

**Supplementary material 2: Cross-links identified within the IFT-B1_B2 hexamer by XL-MS/MS**

Same as for Supplementary material 1 but carried out on the IFTB1_B2 complex (see text and Figure 1).

**Supplementary material 3: List of point mutations in IFT-B associated with human ciliopathies**

The legend is embedded in the excel file.

**Supplementary Table 1:** X-ray data collection and refinement statistics for the CrIFT70/52/88 and CrIFT70/52/46 crystals.

| **Protein complex** | **CrIFT70_52_88** | **CrIFT70_52_46** |
|---|---|---|
| **Wavelength (Å)** | 1.00 | 1.00 |
| **Resolution range (Å)** | 10 - 3.75 (4.02 - 3.75) | 90 - 4.0 (4.24 - 4.0) |
| **Space group** | C 1 2 1 | P 1 21 1 |
| **Unit cell (a,b,c,Å)** | 118,0, 269.5, 95.8, 90.11 | 98.4, 113.4, 374.5, 90.06 |
| **Total reflections** | 85471 (16160) | 476912 (47298) |
| **Unique reflections** | 24628 (4890) | 70497 (10960) |
| **Multiplicity** | 3.4 (3.3) | 6.8 (6.4) |
| **Completeness (%)** | 95.2 (94.7) | 99.4 (91.4) |
| **Mean I/sigma(I)** | 7.8 (1.5) | 7.7 (0.6) |
| **Wilson B-factor (Å2)** | 145 | 184 |
| **Twin law** | NA | h,-k,-l |
| **R-pim** | 0.054 (0.63) | 0.086 (1.6) |
| **CC1/2** | 0.998 (0.626) | 0.972 (0.325) |
| **Reflections used in refinement** | 24628 (3616) | 70497 (10960) |
| **Reflections used for R-free** | 1442 (210) | 2913 (188) |
| **R-work** | 0.358 (0.433) | 0.313 (0.346) |
| **R-free** | 0.379 (0.462) | 0.346 (0.385) |
| **Number of non-hydrogen atoms** | 12261 | 22980 |
| **macromolecules** | 12261 | 22980 |
| **Ligands** | 0 | 0 |
| **Solvent** | 0 | 0 |
| **Protein residues** | 1548 | 2976 |

| | | |
|---|---|---|
| **RMS(bonds)** | 0.004 | 0.006 |
| **RMS(angles)** | 0.77 | 1.32 |
| **Ramachandran favored (%)** | 95.60 | 91.49 |
| **Ramachandran allowed (%)** | 4.20 | 7.92 |
| **Ramachandran outliers (%)** | 0.20 | 0.59 |
| **Rotamer outliers (%)** | 0.84 | 0.04 |
| **Clash score** | 14.85 | 29.9 |
| **Average B-factor (Å$^2$)** | 185 | 198 |

**Supplementary Table 2:** SAXS Data collection parameters for IFT38$_{1-133}$ and IFT80$_{1-582}$/38$_{1-133}$ complex

| | | | |
|---|---|---|---|
| **Data-collection parameters** | | | |
| Instrument: | | ESRF BM29 | |
| Wavelength (Å) | | 0.99 | |
| q-range (Å$^{-1}$) | | 0.0032 – 0.49 | |
| Sample-to-detector distance | | 2.867 m | |
| Exposure time (sec) | | 1 per frame | |
| Temperature (K) | | 283 | |
| Detector | | Pilatus 1M (Dectris) | |
| Flux (photons/s) | | 1 × 10$^{12}$ | |
| Beam size (μm$^2$) | | 172 × 172 | |
| **Structural parametes** | | IFT38$_{1-133}$ | IFT80$_{1-582}$/38$_{1-133}$ |
| Type of experiment | | CS | CS |
| Concentration used (mg/mL) | | 0,4-6,4 (0,4) | 0,37-6 (0,75) |
| NaCl concentration (mM) | | 150 | 150 |
| From p(r) | Rg (Å) | 21,5 | 29,2 |
| | Dmax (Å) | 70,4 | 102,2 |
| Porod volume Vp x10$^3$ (Å3) | | 21,97 | 106,37 |
| Molecular mass (kDa) from Vp | | 13,7 | 66,4 |
| Molecular mass (kDa) from sequence | | 15 | 80 |
| **Modeling** | | | |
| Dammif | NSD | 0,557 | 0,628 |
